# Detecting and dissecting signaling crosstalk via the multilayer network integration of signaling and regulatory interactions

**DOI:** 10.1101/2022.09.29.510183

**Authors:** Arda Halu, Seung Han Baek, Ian Lo, Leonardo Martini, Edwin K. Silverman, Scott T. Weiss, Kimberly R. Glass

## Abstract

The versatility of cellular response arises from the communication, or crosstalk, of signaling pathways in a complex network of signaling and transcriptional regulatory interactions. Understanding the various mechanisms underlying crosstalk on a global scale requires untargeted computational approaches. We present a network-based statistical approach, MuXTalk, that uses high-dimensional edges called multilinks to model the unique ways in which signaling and regulatory interactions can interface. We demonstrate that the signaling-regulatory interface is located primarily in the intermediary region between signaling pathways where crosstalk occurs, and that multilinks can differentiate between distinct signaling-transcriptional mechanisms. Using statistically over-represented multilinks as proxies of crosstalk, we predict crosstalk among 60 signaling pathways, expanding currently available crosstalk databases by more than five-fold. MuXTalk surpasses existing methods in terms of prediction performance, identifies additions to manual curation efforts, and pinpoints potential mediators of crosstalk for each prediction. Moreover, it accommodates the inherent context-dependence of crosstalk, allowing future applications to cell type- and disease-specific crosstalk.

## Introduction

Signal transduction pathways are sequences of biomolecular interactions through which cells respond to their environment, processing internal and external stimuli through a succession of receptors, intermediate signaling proteins, transcription factors (TFs) and their target genes. Signaling pathways do not perform their functions in isolation; they instead operate within a global network of protein-protein interactions (1, 2). Due to the tightly knit and often overlapping way in which signaling pathways are embedded in this global network, stimuli received by one pathway frequently lead to downstream effects in another pathway, resulting in what is called signaling crosstalk. Signaling crosstalk helps to combine the relatively few canonical signaling pathways in different ways, giving rise to the vast range of cellular responses observed in healthy development and homeostasis, as well as in disease (3, 4).

Analyzing signaling networks presents numerous challenges. First, signaling networks consist of multiple types of edges, i.e., signaling interactions. Some types of signaling interactions number in the thousands, whereas other types include only a handful. Second, signaling edges are highly context-dependent: An edge between the same pair of nodes may represent different relationships (e.g., inhibition vs activation) in different pathways. Third, signaling is inherently entangled with transcriptional regulation since TFs and their targets form the last stage of signaling cascades. Indeed, the transcriptional control of a pathway by another pathway has been shown to be an important contributing mechanism to crosstalk (5–7).

The computational modeling of signaling crosstalk has traditionally focused on “critical” nodes, i.e., the common genes between pathways, that were deemed to serve as a junction to cause crosstalk (8, 9). This idea has later been extended to gene sets, inferring crosstalk by the similarity between enriched functional annotation terms (10). Network methods have added more sophistication to these overlap- and enrichment-based approaches by defining crosstalk based on the interactions that lie between signaling pathways (11–15). Recently, further refined network-based approaches that consider the canonical definition of signal transduction have emerged (16, 17). These methods model crosstalk as the propagation of signals in an underlying network from receptors to TFs and consider long-range interactions between signaling pathways.

Despite the utility and individual advantages of each of these approaches, each has one or more of the following limitations that impact their specificity, predictive power and context-specific application: (i) Treating signaling pathways as gene sets and disregarding the interactions of individual genes within and between them; (ii) Not differentiating between the multiple types of signaling interactions; (iii) Focusing solely on the set of receptors and TFs in a pathway as the input and not the remaining proteins in the pathway; (iv) Not modeling additional crosstalk mechanisms such as crosstalk by PPIs, feedback loops, and downstream TF targets that are members of other pathways; (v) Relying on absolute numbers of edges without providing a statistical background; (vi) Being limited in scope, typically restricting their analyses to a handful of the most well-described signaling pathways.

In this study, we posit that a deeper understanding of cellular signaling, including crosstalk, requires a unified view of signal transduction and transcriptional regulatory events in the cell (18, 19). To integrate cell signaling and gene regulation while tackling the above-mentioned challenges around analyzing signaling networks, we use multilayer networks (20, 21), which can simultaneously keep track of multiple types of concurrent and context-dependent edges in the form of distinct layers of networks. In particular, we use high-dimensional edges called *multilinks* (22, 23) to model the unique ways in which signaling and regulatory interactions can interface with each other. We introduce a computational framework, MuXTalk, that exploits the statistics of multilinks to characterize signaling pathways and to study the mechanisms and potential molecular conduits contributing to their crosstalk. In our benchmarks, MuXTalk performs better than gene set-based and other network-based approaches in identifying crosstalk. We find recent literature evidence of crosstalk previously not captured in extensive curation efforts, signifying additions to current benchmarks. Crosstalk predictions in our “discovery” set of pathway pairs are highly supported in the literature. Overall, MuXTalk addresses the limitations outlined above by being a network-based, statistical approach that can accommodate context specificity in the biomolecular interactions at the signaling and transcriptional regulation levels and model multiple crosstalk mechanisms. Covering 60 KEGG signaling pathways, it expands the search space of the currently available crosstalk databases with a more than five-fold increase of potentially crosstalking pathway pairs. The MuXTalk package and web app are available at (https://github.com/r-duh/MuXTalk).

## Results

### The multilayer network of signaling and regulatory interactions

To jointly analyze signal transduction and transcriptional regulatory interactions, we built a multilayer network that consists of a signaling layer and a regulatory layer (Figure 1A). We first constructed a network of known signaling events in human using signaling pathways curated from the KEGG database (24) **(Methods)**. The resulting KEGG signaling network, comprised of 60 signaling pathways (Supplementary Table 1), was a directed multigraph (i.e., a network that can have multiple directed edges between the same pair of nodes) with 2,363 genes and 40,966 edges encompassing 11 interaction types (Figure 1B). Despite being diverse in size, with anywhere from 6 to 294 nodes, most signaling pathways had similar network densities (<0.1) (Figure 1C, Supplementary Table 1). The edges in the KEGG signaling network had high edge multiplicity (i.e., multiple edges between the same pair of nodes) with 18% of edges being present in multiple pathways, typically four or fewer (Supplementary Figure 1A). Furthermore, a sizeable portion (17%) of these edges (635 edges in total, corresponding to 3% of all signaling edges) represented different types of interactions in different pathways (Supplementary Figures 1B-C). As an example, the edge from NF1 to KRAS is inhibitory in the MAPK signaling pathway, whereas the same edge represents activation in the Ras signaling pathway.

**Figure 1.**
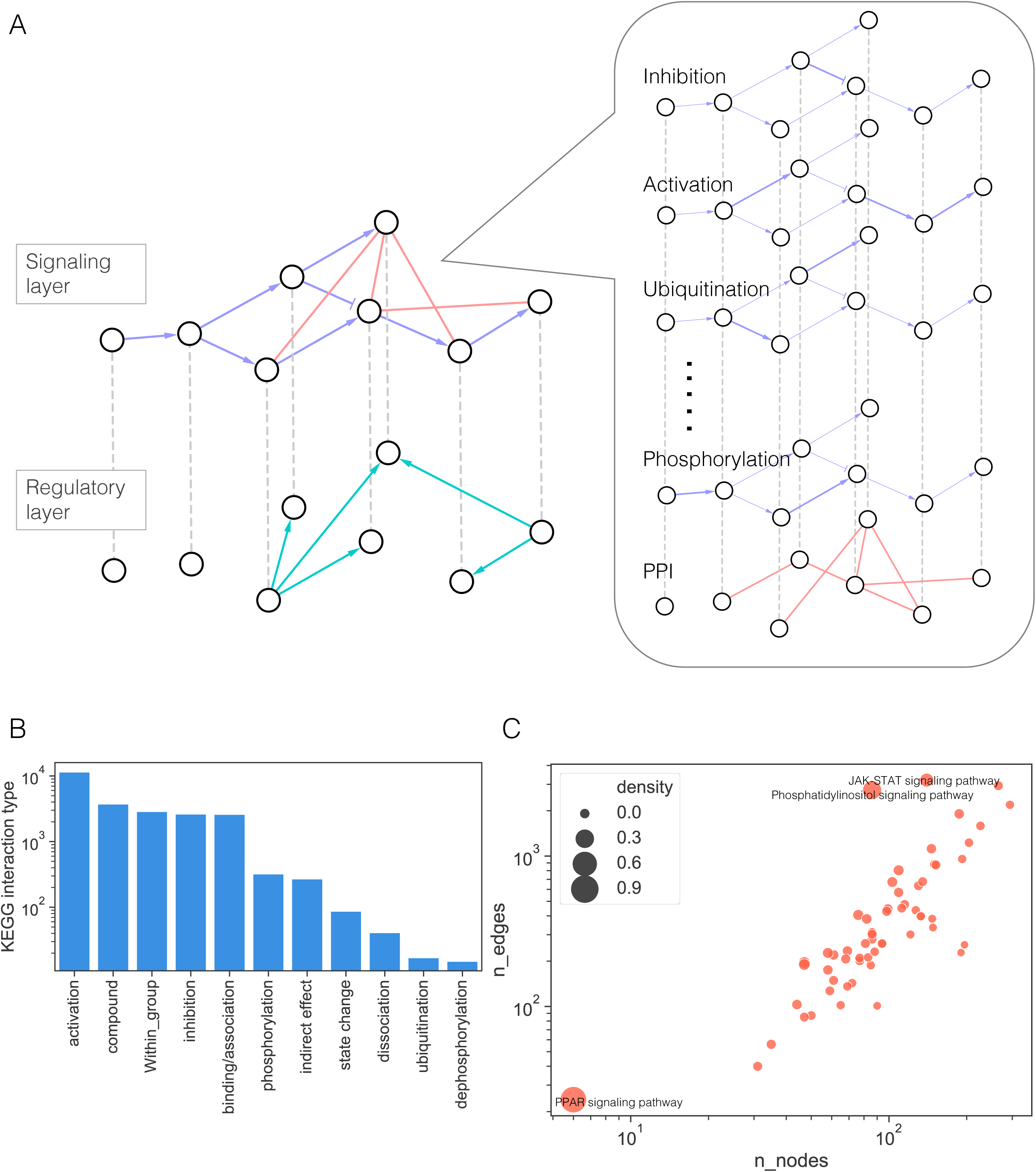
**(A)** Schematic showing the multilayer network consisting of a regulatory layer and a signaling layer, which in turn comprises individual layers for each type of signaling interaction in the KEGG signaling network and a “context-free” protein-protein interaction (PPI) network that acts as a scaffolding for signaling interactions. Due to multiple signaling pathways being superimposed, nodes can be connected by multiple parallel edges in the signaling layer. **(B)** The number of unique edges belonging to each signaling interaction type. **(C)** The number of nodes and edges for each KEGG signaling pathway. Circle sizes correspond to network density, defined by the ratio of the number of existing edges to the number of possible edges given the network size. Pathways with high network density (>0.1) are indicated.

While this directed network of annotated signaling pathways is necessary to provide biological context, it is, by itself, generally not sufficient for discovering new mediating interactions between signaling pathways (19). For this purpose, larger and unannotated (context-free) undirected protein-protein interaction (PPI) networks have often been used as a scaffolding, or “skeleton” network, to underpin efforts to model an organism’s signaling circuitry (1,2,25). We supplemented the KEGG signaling network with a literature-curated, large-scale PPI network (26), whose combination with the KEGG network formed the signaling layer **(Methods)**. The addition of the PPI edges expanded the scope of the signaling network to 16,080 nodes and 239,048 edges. For the transcriptional regulatory layer, we used a previously published large-scale human gene regulatory network (GRN) (27). To ensure the robustness of our approach for networks with different densities, we used GRNs generated using three different p-value thresholds, which resulted in GRN densities spanning three orders of magnitude (Supplementary Table 2) **(Methods)**.

### Signaling-regulatory interface is located in the intermediary network region between signaling pathways

Overlapping signaling and regulatory interactions signify potential points of interface between the signaling and regulatory networks that might contribute to crosstalk (18). To quantify the degree of this interfacing, we measured the overlap between signaling and regulatory edges across the entire multilayer network. We compared the actual number of overlapping edges with the overlap observed between the two layers in randomized networks **(Methods)**. There were 226 edges common to both the signaling and the regulatory layer, which was significantly higher than random expectation (z-score=7.53, empirical p-value<0.002) (Figure 2A). This observation was independent of the density of the regulatory layer (Supplementary Figures 2A, C). In contrast with this overall edge overlap, the signaling and regulatory edge overlap within KEGG signaling pathways was not significant for the majority of pathways (median z-score=-0.23) (Figure 2B). Even with two denser GRNs in which a high layer overlap is expected, only about half of the pathways had significant overlap, with a median z-score of 1.13 and 2.98 (Supplementary Figures 2B, D). In total, a high proportion (83-93%) of signaling-regulatory interaction overlap (Supplementary Figure 3) was located outside of the signaling pathways. Together, these results imply that the significant signaling-regulatory interaction overlap is primarily located in the intermediary network region between the KEGG signaling pathways. Since signaling crosstalk is frequently mediated by the interactions between signaling pathways, these overlapping signaling-regulatory interactions that lie between pathways warrant closer investigation and might shed light on crosstalk mechanisms.

**Figure 2.**
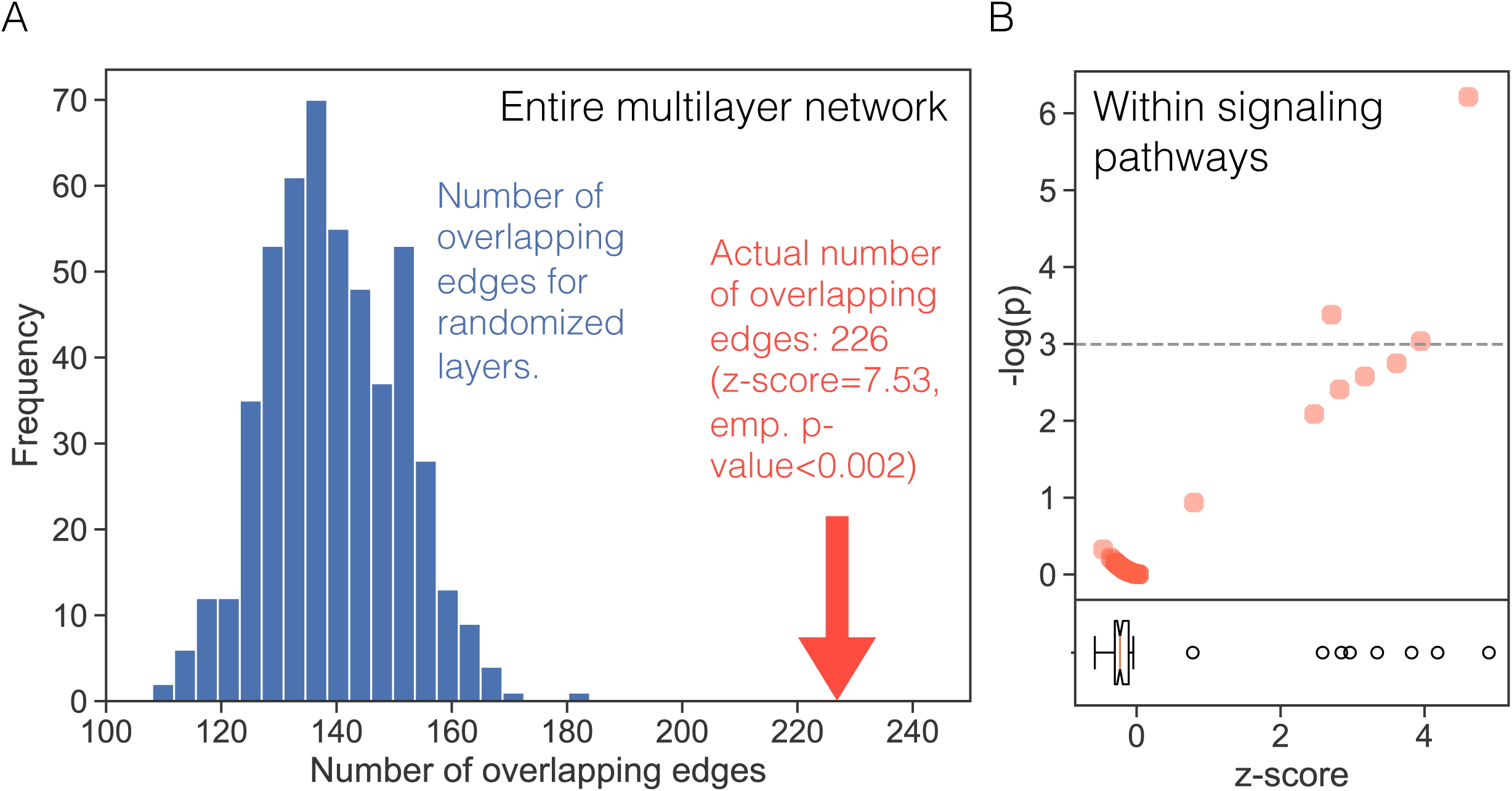
**(A)** The number of overlapping signaling and regulatory edges in the multilayer network with a GRN layer determined using a p-value threshold of p<10^-6^. The blue bars show the distribution of overlapping edges for the randomized networks and the red arrow indicates the overlap for the actual multilayer network. **(B)** The -log(empirical p) values for overlap within each signaling pathway in the multilayer network with the GRN layer (p-value threshold of p<10^-6^). Each dot represents a KEGG pathway. The boxplot indicates the distribution of z-scores for overlap within each pathway.

### Multilink statistics reveal mechanisms between signaling and regulatory interactions

To integrate signaling and regulatory interactions, we rely on high-dimensional edges called *multilinks* (22, 23), which enable us to simultaneously keep track of multiple types of concurrent and context-dependent interactions present in a multilayer network. Concretely, multilinks enumerate every possible combination of interaction types across all layers of the multilayer network as a unique type of edge, each of which exemplifies a distinct signaling/regulatory mechanism (Figure 3A). We hypothesized that the relative abundance of multilink types can be used to assess the contribution of each mechanism to a given signaling pathway and to subsequently predict crosstalk between signaling pathways. We developed a statistical framework, named MuXTalk, that (i) determines whether each multilink type is over- or under-represented in the multilayer network, and (ii) identifies potentially crosstalking pairs of pathways using a certain class of multilinks as a proxy of crosstalk. Briefly, MuXTalk performs the following steps (Figure 3B-C): (i) It counts the number of occurrences of each multilink type in the multilayer network; (ii) It compares these numbers to those obtained from a large ensemble of degree-preserved randomized networks to calculate statistics (z-scores and empirical p-values) associated with each multilink type; (iii) It prioritizes pairs of pathways as the most likely to crosstalk using the multilink statistics of the edges between signaling pathways **(Methods)**.

**Figure 3.**
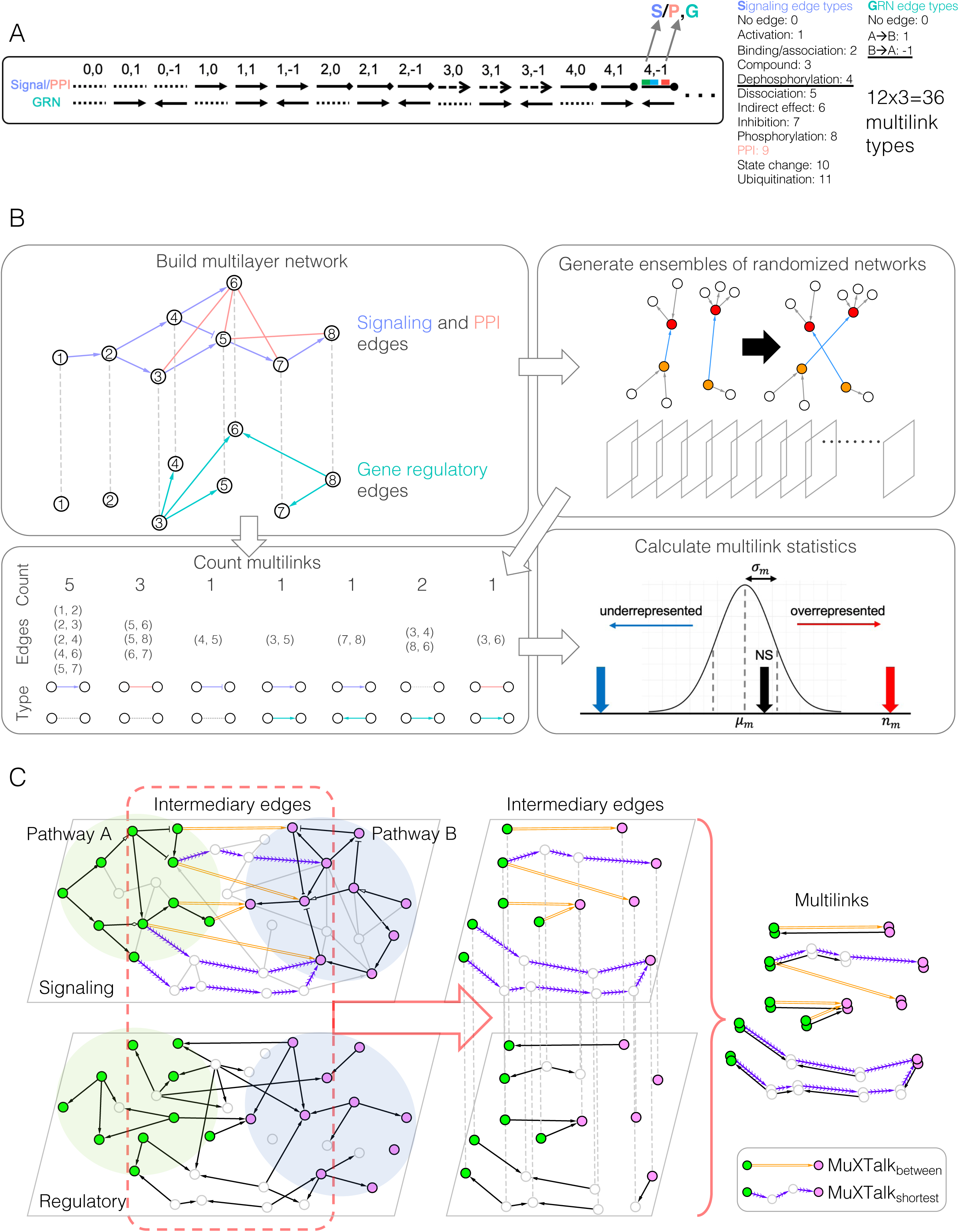
**(A)** Overview of multilink types (*S, R*) between the signaling and regulatory layers where *S* ∈ [0, 11] and *R* ∈ {−1, 0, 1} are integers representing signaling and regulatory edge types, resulting in a total of 36 multilink types. **(B)** Overview of the MuXTalk framework. **(C)** Description of MuXTalk_between_ and MuXTalk_shortest_, which identify crosstalk based on the multilinks of direct edges and shortest paths between pairs of pathways, respectively.

For a more detailed picture of signaling and regulatory overlap across all KEGG signaling pathways, we first used MuXTalk to calculate the multilink statistics of all edges in the multilayer network (Figure 4). Overall, multilinks of type (0, 1) and (0, −1), which represent the absence of an edge in the signaling layer and the presence of an edge in the regulatory layer (Figure 3A), were significantly under-represented across all signaling pathways, supporting our finding of significant overlap between the two layers in the previous section. When broken down into individual signaling events, dephosphorylation, dissociation, inhibition, phosphorylation and protein-protein interactions were often accompanied by regulatory interactions. Dephosphorylation, inhibition and phosphorylation events were in opposite direction with regulatory events (i.e., (4, −1), (7, −1) and (8, −1) multilink types were significantly over-represented), whereas dissociation events were in the same direction as regulatory events (i.e., (5, 1) was significantly over-represented). In contrast, activation, binding/association, and compound interactions tended to occur without concurrent regulatory interactions (i.e., multilink types (1, 0), (2, 0) and (3, 0) were over-represented. These results are concordant with signaling-regulatory mechanisms documented in the literature: Phosphorylation and dephosphorylation can directly regulate transcription factor function (28, 29); inhibition of certain pathways has been shown to antagonize transcription factor function (30); the dynamics of dissociation of TFs from their targets have been shown to determine their function (31); activators tend to be co-activator TFs and are therefore generally not targets of TFs.

**Figure 4.**
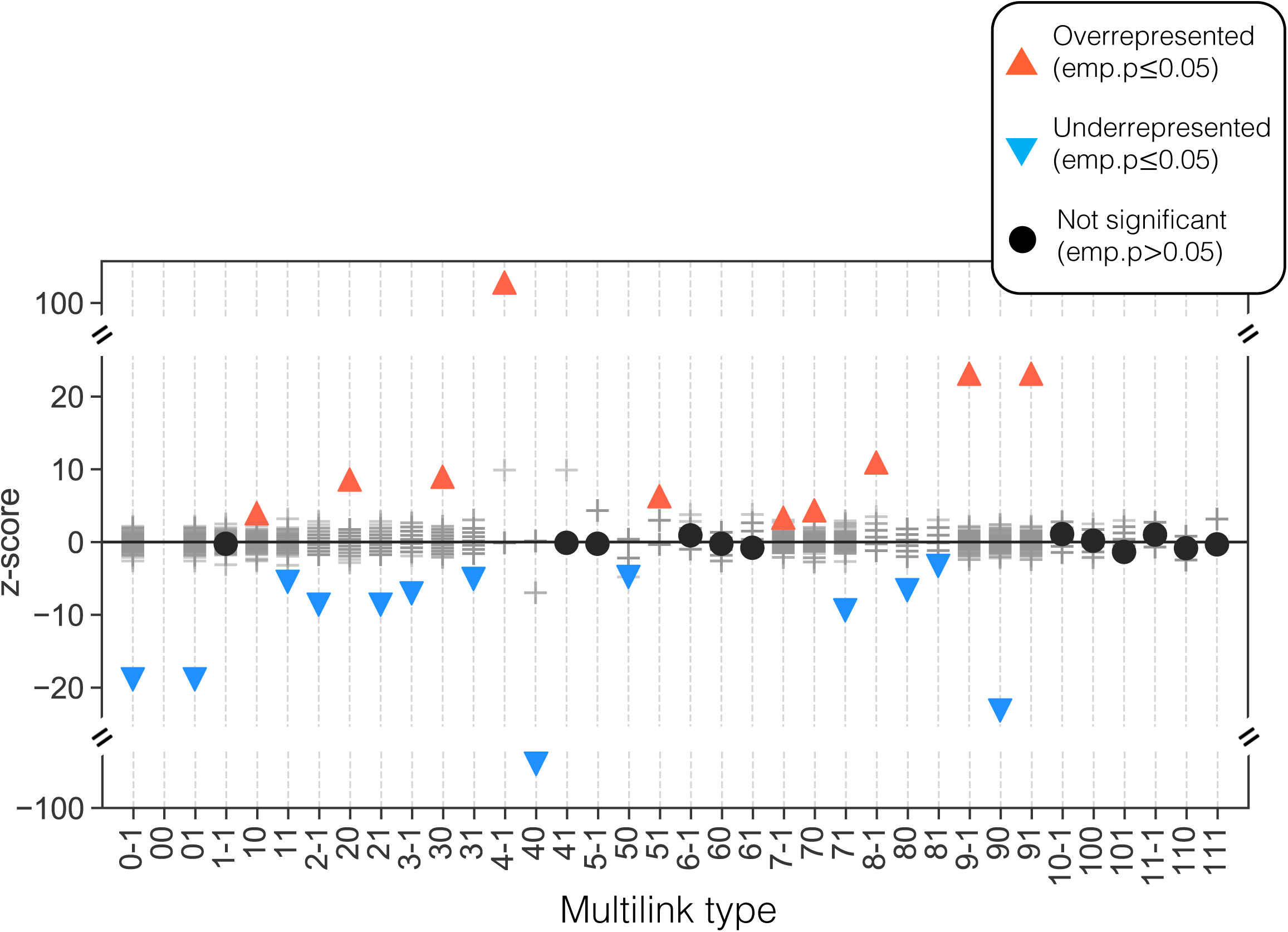
Multilink profiles of all edges in the multilayer network with a GRN layer based on a p-value threshold of p<10^-4^. Multilink types are represented by numerical indicators where the first one or two digits represent the signaling edge type and the last digit represents the regulatory interaction direction. Red triangles pointing up and blue triangles pointing down indicate statistically over-represented (z>0; emp. p≤0.05) and under-represented (z<0; emp. p≤0.05) multilink types, respectively. Black circles denote statistically insignificant multilink types (emp. p>0.05).

Notably, the over- and under-represented multilink types were consistent across GRNs through the entire range of densities tested, with the exception of some missing multilinks due to the limitations of calculating multilink statistics in sparser GRNs (Supplementary Figure 4). Further, the significantly over- and under-represented multilinks within individual signaling pathways were highly pathway-specific, showing high variability in terms of which mechanisms were predominantly featured in each pathway, including more “elusive” mechanisms such as ubiquitination and state change (Supplementary Figures 5-7). Together, these results suggest that multilink statistics can be a useful tool in extracting signaling-regulatory mechanisms within and between signaling pathways, offering clues as to which mechanisms may be involved in signaling crosstalk.

### MuXTalk outperforms other methods in predicting crosstalk

After exploring the usefulness of multilinks in elucidating signaling-regulatory mechanisms, we next tested our hypothesis that multilink statistics of edges in the intermediary region between signaling pathways can be harnessed to identify their crosstalk. Since crosstalk can occur when signaling pathways connect either directly or indirectly (3), we implemented two complementary approaches within MuXTalk, namely, MuXTalk_between_ and MuXTalk_shortest_ (Figure 3C), which consider the direct edges and long-range interactions (i.e., shortest paths) between pathways, respectively **(Methods)**. Using the over-representation of multilink types that include regulatory edges (i.e. multilink types (*S*, 1) and (*S*, −1), see **Methods**) as a proxy of crosstalk, we can capture, at once, multiple mechanisms through which crosstalk can occur, including directed signaling interactions and undirected protein-protein interactions, as well as regulatory mechanisms such as transcription factors of a pathway targeting members of another pathway (32, 33) and ensuing feedback loops (34–36).

To assess the validity of the MuXTalk predictions and compare the prediction performance to those of other methods, we designed two benchmarks, one deterministic and one stochastic, based on crosstalking pathway pairs from the literature-curated crosstalk database XTalkDB (37) **(Methods)**. Since XTalkDB only includes 25 of the 60 KEGG signaling pathways considered here, we used these 25 pathways as our “benchmark set” and the remaining pathways as our “discovery set” (Supplementary Table 1). As exemplary methods to compare MuXTalk with, we chose four other network-based approaches that rely on node and edge overlap (8, 9), direct edges (13, 38) and shortest paths (16) between pathways. Both versions of MuXTalk surpassed all other methods in terms of the area under both the receiver operating characteristic (ROC) curve and the precision-recall (PR) curve, in both the deterministic and the stochastic version of the benchmark (Figure 5A, Supplementary Figure 8). These results were recapitulated when we used denser GRNs (Supplementary Figure 9), suggesting the robustness of MuXTalk against changes in network density. Despite a slight drop in the performance with the densest GRN, we see its sustained advantage over alternative methods (Supplementary Figure 9).

**Figure 5.**
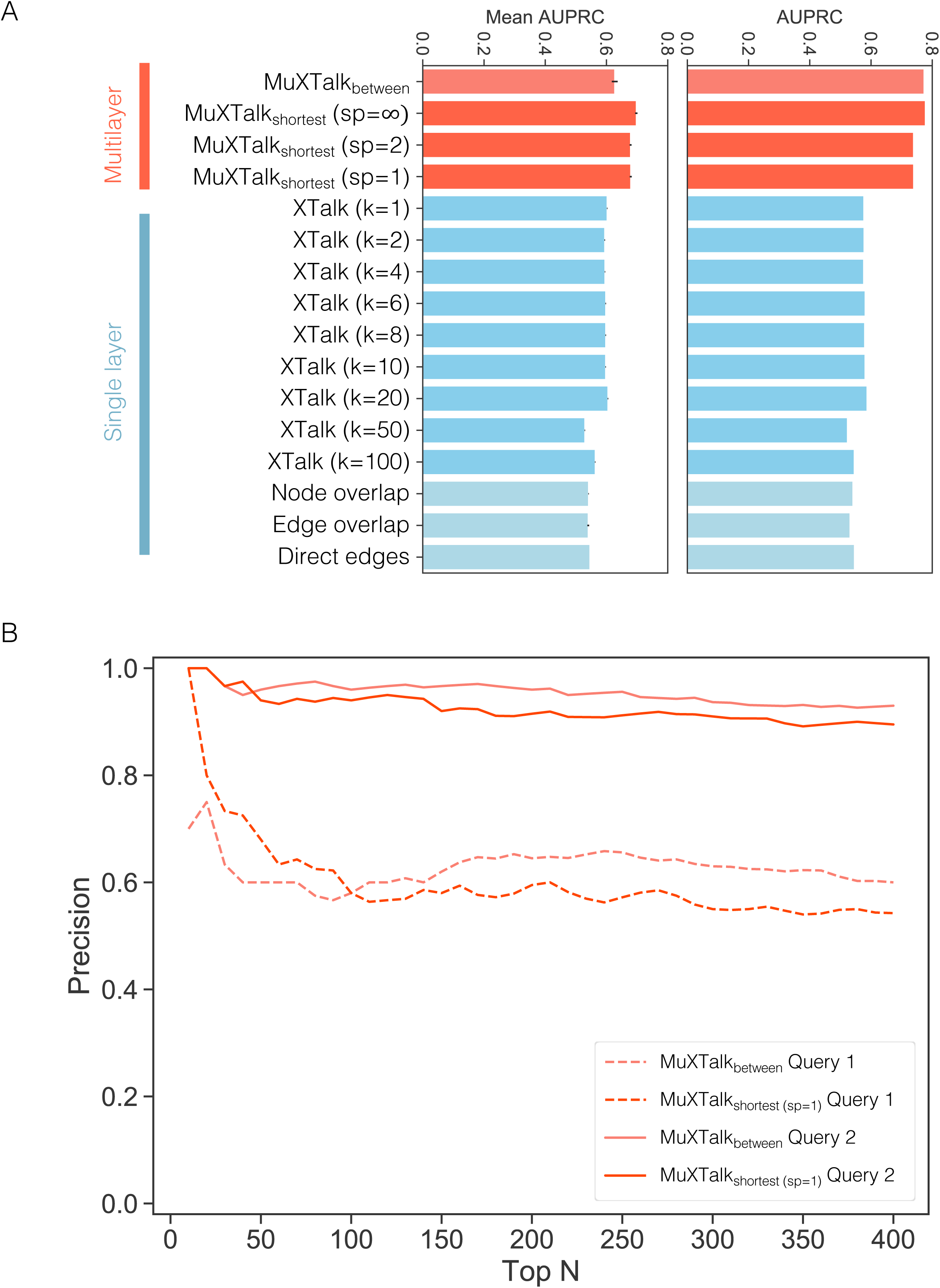
**(A)** Area under the precision-recall curves (AUPRC) for MuXTalk (red) and four other methods (blue) for the stochastic (left) and deterministic (right) versions of the benchmark. Error bars indicate the standard deviation. MuXTalk was run on the multilayer network using a GRN layer based on a p-value threshold of p<10^-6^ **(B)** Precision-rank plot for the top 400 predictions in the discovery set of pathways. Dashed and solid lines are for PubMed Query 1 and 2, respectively.

### Discovering new putative crosstalk connections with MuXTalk

Encouraged by the better performance of MuXTalk compared to other network-based methods, we delved further into MuXTalk’s predictions in the benchmark set. Since the benchmark data (37) is several years old (published in 2016), we queried PubMed to look for recent publications that might provide additional evidence that could further support MuXTalk’s predictions. We performed two separate PubMed queries to capture potential crosstalk among the 600 possible pathway pairs in the benchmark set **(Methods)**. Overall, 42 putative crosstalking pathway pairs detected by MuXTalk_shortest_ were not in our benchmark set in either direction. 21 of these pairs had PubMed literature evidence by Query 1 and all of them had PubMed literature evidence by Query 2 (Supplementary Table 4). Similarly, 13 putative crosstalking pathway pairs detected by MuXTalk_between_ were not in our benchmark set in either direction; 4 and 12 of these had PubMed literature evidence by Query 1 and Query 2, respectively (Supplementary Table 5). Some of these literature hits were from after 2016, suggesting the utility of MuXTalk in supporting manual curation efforts with its predictions. We next used MuXTalk on the discovery set of pathway pairs to identify potential signaling crosstalk between pathways not included in the benchmark set. By Query 2, both MuXTalk_shortest_ and MuXTalk_between_ showed a mean precision of 98% for the top 50 predictions. In terms of Query 1, MuXTalk_shortest_ and MuXTalk_between_ had a mean precision of 79% and 66%, respectively, for the top 50 predictions (Figure 5B). Together, these results provide support for the novel crosstalk predictions made by MuXTalk, as well as represent potential additions to the current crosstalk databases.

### MuXTalk sheds light on crosstalk mechanisms; identifies potentially novel mediators of crosstalk

To demonstrate MuXTalk’s potential utility for discovering novel crosstalk events and providing mechanistic insights to each, we focused on the highest ranked crosstalking pairs identified by MuXTalk. Both MuXTalk_between_ and MuXTalk_shortest (sp=1)_ identified the crosstalk between the neurotrophin signaling pathway and the TGF-β signaling pathway among the top-ranked crosstalking pathway pairs previously not documented in the benchmark data (Supplementary Tables 4 and 5).

While the crosstalk from the TGF-β signaling pathway to the neurotrophin signaling pathway, as captured by MuXTalk_between_, has been previously established due to the transcriptional regulatory role of TGFβ signaling on neurotrophins (39–41) the crosstalk in the opposite direction has been relatively underappreciated until recently (42) wherein it was speculated that neurotrophins and TGF-βs act in concert to activate a mutual protective signaling network. Supporting this notion, MuXTalk_shortest_ discovered two statistically significant multilinks of type (9, 1) (meaning a PPI and a regulatory interaction, see Figure 3A) pointing from the neurotrophin signaling pathway to the TGF-β signaling pathway (Figure 6), connecting Activating Transcription Factor 4 *(ATF4)* with DNA Damage Inducible Transcript 3 *(DDIT3)* and Tumor Protein P73 *(TP73)* with Cyclin G1 *(CCNG1)*. *DDIT3* and *CCGN1* here serve as the intermediary molecules that interact with E1A Binding Protein P300 *(EP300)* and Cullin 1 *(CUL1)* and Protein Phosphatase 2 Catalytic Subunit Alpha *(PPP2CA)* and Protein Phosphatase 2 Scaffold Subunit Aalpha *(PPP2R1A)*, respectively, in the TGF-β signaling pathway.

**Figure 6.**
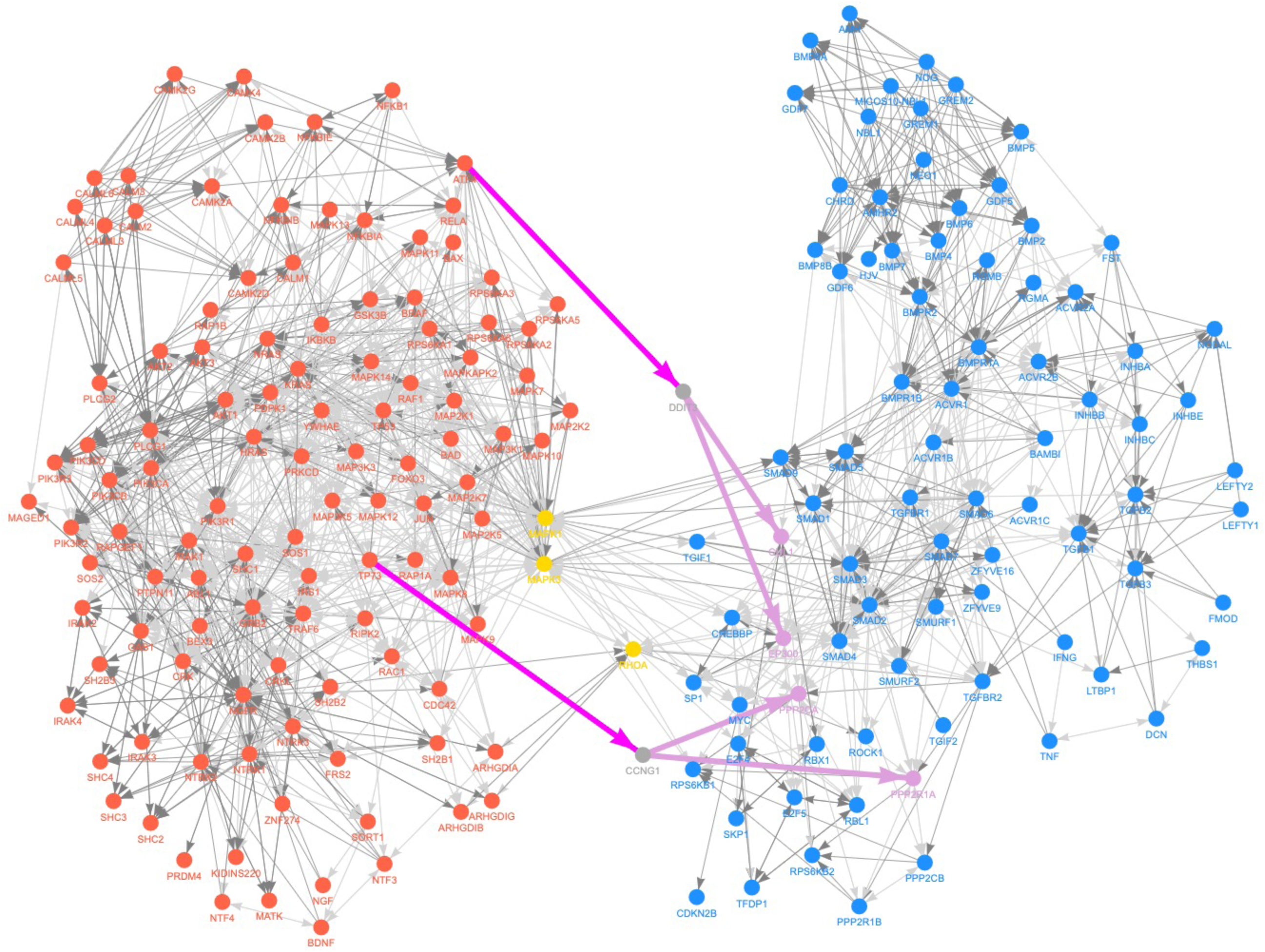
Example output from the MuXTalk web app that shows the crosstalk from the neurotrophin signaling pathway (red) to the TGF-β signaling pathway (blue), predicted by MuXTalk_shortest_ (with sp=1). Orange nodes indicate the genes common to both pathways. Purple edges indicate significant multilinks of type (9, −1) (PPI edge and a GRN edge from the neurotrophin signaling pathway to the TGF-β signaling pathway) that were identified by MuXTalk to mediate the crosstalk.

The impact of *DDIT3* (also known as the growth arrest and DNA damage-inducible protein 153 *(GADD 153)* and encodes the C/EBP homologous protein (*CHOP*)) in diverse human diseases including neurodegenerative disorders has been established, whereas the emerging role of *DDIT3* in fibrosis has only recently been reported (43). In particular, the crucial role of *DDIT3*/*CHOP* expression in the negative regulation of two neurotrophic cytokines, leptin and insulin-like growth factor-1, has been reported (44), whereas other mechanistic studies showed that *CHOP* regulates the production of pro-inflammatory (M2) macrophages and subsequent TGF-β signaling involved in idiopathic pulmonary fibrosis (IPF) (45). In turn, the inhibition of the two interactions of *DDIT3* identified by MuXTalk, *CUL1* and *EP300*, has been shown to reduce fibroblast proliferation in chronic obstructive pulmonary disease (COPD) and IPF (46, 47).

TP73 is a member of the p53 family of transcription factors that targets neurotrophin receptor (p75NTR), promoting terminal neuronal differentiation (48, 49). Neurotrophin signaling mediated through p75NTR has been shown to activate the nuclear factor-kB (NF-kB) and Jun kinase signaling pathways (including JNK pathway) (50, 51). While the NF-kB and JNK pathways play an important role in neural development in many aspects, it is also known that they regulate the expression and activation of cyclins, including Cyclin G1 (CCNG1) (52–54). More specifically, the NF-kB signaling pathway was shown to regulate Cyclin D1 (CCND1) expression, whereas the JNK signaling pathway was shown to regulate the expression of both CCND1 and CCNG1 (52–54). CCNG1 is a non-canonical cyclin that was initially discovered as a novel member of the cyclin family (55). Although the exact mechanism is not yet clear, CCNG1 is known to play a crucial role in cell proliferation and cell growth (56–58). Indeed, CCNG1 was shown to directly interact with Protein Phosphatase 2A (PP2A), which is a holoenzyme complex that consists of the PPP2CA and PPP2R1A subunits, and recruits PP2A to MDM2 (55,59–62). The PP2A mediated de-phosphorylation and activation of MDM2, results in the destabilization of P53, which is crucial in cell proliferation, signal transduction and apoptosis (55,61,62). While PP2A may influence TGF-β signaling through P53 indirectly, PP2A was also shown to directly regulate TGF-β signaling by de-phosphorylating the TGF-β receptor (63, 64).

## Discussion

Signaling pathways combine with and modulate each other in myriad ways in an intricate web of signaling and regulatory interactions, allowing cells to fine-tune their responses to their microenvironment. Canonical signaling pathways and their crosstalk have long been subjects of targeted studies that derive mechanistic insights in the context of a disease or developmental process of interest. Given the combinatorial space within which signaling pathways can interact, both at the physical protein interaction and transcriptional regulation level, untargeted computational methods can provide a holistic view of crosstalk by analyzing the global network of signaling interactions and offering putative crosstalk mechanisms in a context-specific way. In this study, we introduced a multilayer network-based statistical framework to integrate signaling and regulatory interactions, which allowed us to predict crosstalk events and identify their potential mediators.

Dedicated manual curation efforts have resulted in highly reliable crosstalk databases such as XTalkDB that continue to inform computational efforts (16), including ours. Still, by definition, such studies rely on existing knowledge: They use as their starting point keyword searches in PubMed and then filter these automated results based on expert curation. As such, they have the potential to miss the subtler ways in which signaling crosstalk can be described in the literature. Our approach, by contrast, takes the molecular bases of crosstalk as the starting point and predicts the potential mediators of crosstalk, potentially transcending the limitations of what is already published in the literature and uncovering novel mechanistic insights. Indeed, many of the studies supporting our use cases have been published after these crosstalk databases were established, suggesting the potential complementary role of “prospective” computational models to “retrospective” literature curation.

Statistical methods that can distinguish meaningful signals from the noise rampant in biomedical data (65, 66) ought to be preferred where possible. In the context of biological networks, combating noise translates into crafting null network models that generate relevant controls (67–69). When we are interested in the edges that connect signaling pathways as proxies of crosstalk, a highly pertinent question to ask is whether the observed number of connecting edges is more or less than what would be expected by chance, given the connectivity structure of each network. The recently published version of SignaLink (70), which identifies signaling crosstalk and is similar to our work in that it is integrative and offers context-specificity, differs from our method in this important aspect: It is not a statistical framework; it does not perform any hypothesis testing but, rather, simply chronicles the number of connecting edges, and hence does not provide a sense of whether the number of edges between a pair of pathways is statistically meaningful within the underlying network of interactions. With MuXTalk, we generate interaction type-specific ensembles of randomized networks that act as null models and provide a standardized backdrop against which the number of edges can be compared, ensuring that the statistically over-represented multilinks are not simply byproducts of implicit and systematic data biases (71, 72).

One of the limitations of our approach is that it fundamentally relies on knowledge repositories where genes and their products are organized into canonical pathways according to their functional annotations, which themselves might be incomplete (73). There are several such pathway databases with complementary strengths, although traditionally, unifying them has been deemed nontrivial due to varying standard formats and data models (74). As a result, the choice of which database to employ is often based on the specifics of each use case. We chose KEGG for our framework since it is widely used and recognized, has high granularity in terms of identifying distinct interaction types, and can be directly harmonized with XTalkDB for benchmarking. Although it is certain that database choice has at least some bearing on downstream analyses, there is emerging evidence that the disparities between pathway resources might be less consequential than previously estimated: Pathways across different databases were recently found to display much higher levels of agreement when database structural differences were corrected for (75).

Computational constraints generally force a tradeoff between detailed yet small-scale models that simulate signaling dynamics under different perturbations and large-scale yet static models that describe signaling pathways in terms of their topological features. As an example of the latter class of approaches, our method has the inherent limitation of not accounting for the dynamics and differing timescales of signaling and regulatory events. What MuXTalk lacks in this aspect, it makes up for in terms of the size of the multilayer network, spanning multiple types of interactions between over 16,000 proteins and 60 pathways. The study of crosstalk, in particular, is more amenable to this kind of a large-scale network-based approach since it involves interactions among pathways in the overall signaling network. That being said, scalable dynamic models, both fine-grained (atomistic) and coarse-grained (Boolean), still are indispensable vehicles to understand signaling networks and their crosstalk mechanistically. Several advancements in this area have been made in the past decade: Atomistic motif-based models of crosstalk have been proposed (76) and constraint-based stoichiometric approaches akin to the ones used in metabolic networks have been developed for signaling networks (77). Even though these methods are typically applied on individual signaling pathways, the near future may see the fusing of such dynamical models with global approaches like ours.

MuXTalk supports, by design, the discovery of context-specific crosstalk due to its incorporation of GRNs. The MuXTalk framework allows the input of custom GRNs, and we encourage users to explore this direction with GRNs derived from a certain tissue, cell type or disease, or even with condition-specific GRNs inferred from single-cell expression data (78). However, dedicated benchmarks would be necessary to establish the context-specific use of our approach. It is currently challenging to develop benchmarking strategies for cell type- or disease-specific crosstalk due to the lack of ground truth repositories for context-specific crosstalk. This will undoubtedly become a promising future direction as advances in text mining and natural language processing for biomedical literature (79) gradually allow for context-specific knowledge bases for crosstalk.

## Methods

### Building the signaling layer

We curated a list of 60 cellular signaling pathways from KEGG and downloaded their schemas as KGML (KEGG Markup Language) files in May and October 2020 (see Supplementary Table 1 for a list of pathways). We parsed each KGML file to build a directed graph consisting of KEGG “entries” (nodes) and “relations” (edges). Since KEGG relations include both directed and undirected interaction types, we represented undirected interactions as bidirectional edges. Furthermore, in KEGG pathway maps, composite objects such as protein complexes are often represented as a single entry. We disaggregated such gene groups and protein complexes into their individual genes and represented these gene groups as complete graphs in which every constituent gene was connected to every other gene in the group. To disambiguate these interconnecting edges within gene groups, we separately annotated them as “within_group” edges, in addition to the existing KEGG relations. We aggregated the 60 KEGG graphs into a directed heterogeneous multigraph where each directed edge is annotated with the interaction type as well as the signaling pathway it belongs to, allowing for multiple edges between the same pair of nodes. Overall, this precursor KEGG network had 3,306 nodes and 44,286 edges spanning 4 node types (gene, compound, map, and ortholog) and 13 interaction types (activation, binding/association, compound, dephosphorylation, dissociation, expression, indirect effect, inhibition, phosphorylation, repression, state change, ubiquitination, within group). To build the overall signaling layer, we combined the KEGG signaling network with the large-scale human protein-protein interaction (PPI) network curated by Cheng et al. (26). We represented PPI edges as bidirectional and annotated them separately with the “ppi” edge type. During the merging of the KEGG and PPI networks, if a known signaling interaction from the KEGG network coincided with a PPI edge, the KEGG signaling edge superseded the PPI edge. As part of our quality control procedure, the few KEGG edges with unknown interaction type, as well as the self-loops in the PPI and KEGG networks, were removed from the signaling layer. Finally, we removed the “GErel” type of edges (“expression” and “repression”) from the signaling layer to prevent redundancy with the gene regulatory network layer. After these refinements, the final KEGG signaling network consisted of 2,363 genes and 40,966 edges (24,222 unique edges) spanning 60 signaling pathways and 11 types of signaling interactions.

### Building the signaling-regulatory multilayer network

As the basis of our multilink-based crosstalk prediction framework, we built a multilayer network consisting of a signaling layer and a gene regulatory layer. As the gene regulatory layer, we used the human gene regulatory network (GRN) previously described in (27). Briefly, the human GRN was constructed in (27) by scanning the entire hg19 genome for 695 human TF motifs, and a TF was connected to a gene if the motif hit for that TF was in the promoter region of that gene. We considered in our analyses GRNs created using three motif scan p-value thresholds (p<1×10^-4^, p<1×10^-5^, and p<1×10^-6^) corresponding to a wide range of network densities (Supplementary Table 2). Since our multilayer network is technically a multiplex network in which the complete set of nodes is present in all layers, even if as isolated nodes, we only retained genes and their interactions in the signaling network, discarding other KEGG entries such as “maps,” “orthologs” or “compounds.” We then pruned the GRN to only include the genes present in the signaling layer. The number of TFs, targets, edges, and network densities post-pruning are shown in Supplementary Table 2. We used all three versions of GRNs in our sensitivity analyses (Supplementary Figures 2-7 and 9) and used the p<1×10^-6^ cutoff GRN in the rest of the analyses throughout the manuscript. For memory efficiency in downstream calculations, we stored each layer as a sparse matrix. Furthermore, due to the multigraph nature of the signaling layer, where more than one type of edge can exist between the same pair of nodes, we stored each signaling interaction type *e* as a separate sparse matrix. For a larger rewiring space, we added PPI edges to each interaction-specific KEGG layer (named “KEGG_e” layers). In addition, to avoid multiple-counting of the (0, *R*) type of multilinks (detailed in the following section), we collapsed all KEGG interaction types and PPIs on a single network separately (named the “KEGGPPI” layer). KEGGPPI thus represents all the interactions on the signaling layer in a collapsed form without differentiating between the individual types of signaling interactions or PPIs. Altogether, this resulted in sparse matrices for the following layers: 1 for the GRN layer, 11 for the interaction-specific KEGG_e layers, and 1 for the collapsed KEGGPPI layer (Supplementary Figure 10).

### Counting multilinks and obtaining their statistics

Our crosstalk prediction method relies on the statistics of multilinks, which are higher order edges that can represent unique combinations of multiple types of interactions across multiple network layers in a compact way. As a shorthand for denoting multilinks, we represent signaling interactions by integers *S* in the interval [0, 11], and regulatory interactions by the integers *R* in {−1, 0, 1} to account for directionality, where 0 denotes no edge. Together, this resulted in 12 × 3 = 36 multilink types denoted by the pair of integers (*S, R*). We extracted the number of each multilink type by counting all instances of a given (*S, R*) pair across the tensor formed by the stacked adjacency matrices of the multilayer network. Since this operation quickly becomes computationally burdensome for large multidimensional arrays such as ours (2 × 16080 × 16080), we introduced a custom hash to represent each of the 36 multilink types as a unique number. To correctly account for the concurrent edges in the signaling network, we performed the counting step separately on the multilayer formed by the sparse matrix of each signaling interaction type *S* (Supplementary Figure 10). To prevent multiple-counting, we counted the (0, *R*) type of multilinks, i.e., cases where no edge is present in the signaling layer, on the multilayer network formed by the KEGGPPI layer and the gene regulatory layer.

To assess whether a given multilink count is statistically more or less than what would be expected by chance, we generated ensembles of randomized networks. To counter degree bias, we used a degree-preserving randomization method that relies on the random pairwise rewiring of edges (80). To break potential existing degree correlations between layers, we performed this randomization procedure for each layer separately. We generated and stored 500 randomized networks for each of the 13 layers described above, resulting in 250,000 unique randomized multilayer networks for each signaling type-GRN layer pair. To quantify the extent of over- or under-representation of a given multilink count and its statistical significance, we calculated z-scores and two-tailed empirical p-values, respectively, based on the actual *c_a_* and random counts *c_r_* such that

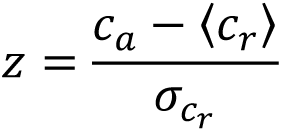

and

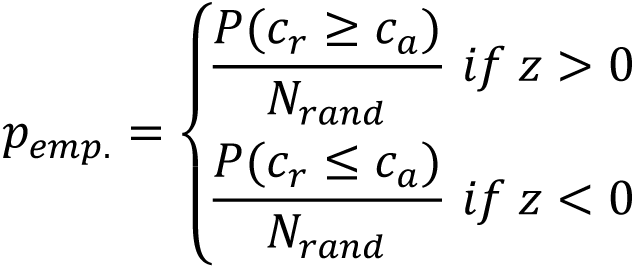

where 〈*c_r_*〉 and *σ_r_* are the mean and standard deviation of *c_r_* and *N*_*rand*_ is the number of random instances. In our analyses, we used *N*_*rand*_ = 100, but this parameter is adjustable by the user. For the reproducibility of our results, the randomized layers used in the benchmarks were drawn in a deterministic manner from this precomputed ensemble (see Supplementary Table 3 for the allocation schema of randomized networks).

### Using multilink statistics to predict crosstalk

In our multilink-based crosstalk prediction framework, we used multilinks of type (*S*, 1) and (*S*, −1) as proxies of crosstalk and deemed as crosstalking the pairs of pathways for which at least one of these multilink types is significantly over-represented (*p*_*emp.*_ ≤ 0.05, *z* > 0). We then devised two approaches to model signaling crosstalk based on multilink statistics: (1) based on the direct edges between a pair of signaling pathways (“MuXTalk_between_”) and (2) based on the shortest paths between a pair of signaling pathways (“MuXTalk_shortest_”) in the signaling layer (“KEGGPPI”).

For MuXTalk_between_, multilink statistics were obtained for the edges connecting Pathway A and Pathway B such that all edges directly connecting the genes in the two pathways were accounted for, excluding (i) the edges within pathways and (ii) the edges connecting the nodes that are common to both pathways (Figure 3C). These criteria were to ensure that only the edges directly between the two pathways were counted. Effectively, we did this by slicing the adjacency matrices of each layer by the mutually exclusive set of nodes in each pathway. We calculated multilink statistics on this final subset of the adjacency tensor.

For MuXTalk_shortest_, multilink statistics were obtained for the edges belonging to the shortest paths connecting Pathway A and B, following the steps below:

i. We determined the mutually exclusive sets of nodes between signaling Pathway A and B.
ii. We for each pair of nodes in this mutually exclusive set of nodes, we computed the shortest path, when such a path exists.
iii. We excluded from these shortest paths any node that belongs to Pathway A or B to identify the set of “intermediary nodes” that exclude the “within-pathway” edges. To fine-tune the reach of the shortest paths between pathways, we introduced a shortest path (sp) threshold that controls the number of intermediary nodes between pairs of pathways. In our simulations, we used sp threshold values of 1, 2 and “no threshold,” meaning that shortest paths of any length were considered.
iv. Using the intermediary nodes, we identified the intermediary shortest paths connecting each pair of mutually exclusive nodes in Pathway A and B.
v. We aggregated all edges in intermediary shortest paths for all sets of mutually exclusive nodes in Pathway A and B. We calculated multilink statistics on this final set of edges.

In both MuXTalk_between_ and MuXTalk_shortest_, we performed the same operations on the randomized multilayer network ensembles and calculated the z-scores and empirical p-values for each multilink type, as described in the previous section. Using these statistics, we then ranked signaling pathway pairs by their (i) number of significantly over-represented (*p*_*emp.*_ ≤ 0.05, *z* > 0) multilink types, (ii) lowest p-value, and (iii) highest z-score. The signaling pathway pairs with the highest number of significantly over-represented multilinks of type (*S*, 1) and (*S*, −1), with the lowest empirical p-values, and with the highest z-scores were thus prioritized as the most likely pathway pairs to crosstalk. To reflect this ranking, we calculated the MuXTalk score as:

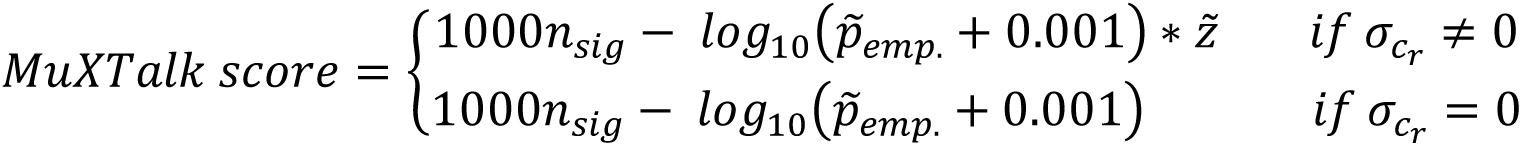

where *n*_*sig*_ is the number of significant multilink types and *p̃*_emp_. and *z̃* are the best empirical p-values and z-scores, respectively.

### Benchmarking

To assess the predictive performance of MuXTalk in comparison with other methods, we devised a benchmark that uses the literature-curated crosstalking pathways available in the XTalkDB database (37). Of the 26 KEGG signaling pathways in XTalkDB, 25 were within the list of 60 KEGG pathways we used to build the signaling layer. Of the 600 possible ordered pairs in these 25 pathways, 331 were marked as crosstalking in XTalkDB.

We compared MuXTalk with node and edge overlap (8, 9), direct edges (13, 38) and shortest paths (16) between signaling pathways. We detail the implementation of each method below:

### Node overlap between signaling pathways

We performed a two-tailed Fisher Exact test to assess the significance of overlap between the nodes of Pathway A and Pathway B given the number of nodes in each pathway and the total number of genes in the 60 KEGG pathways considered. We adjusted the Fisher’s Exact p-values for multiple testing using the Benjamini-Hochberg procedure. We then ranked pathway pairs based on the false discovery rate (FDR) values, with the lowest FDR pathway pair ranking highest as the most likely to crosstalk.

### Edge overlap between signaling pathways

Similar to node overlap, we performed a two-tailed Fisher’s Exact test to assess the significance of overlap between the edges of Pathway A and Pathway B given the number of edges in each pathway and the total number of possible edges in the 60 KEGG pathways considered. We adjusted the Fisher’s Exact p-values for multiple testing using the Benjamini-Hochberg procedure. We ranked pathway pairs based on the false discovery rate (FDR) values, with the lowest FDR pathway pair ranking highest as the most likely to crosstalk.

### Direct edges between signaling pathways

We used direct interactions between pairs of pathways as proxies of signaling crosstalk, as done previously by Korcsmaros et al. (13). For each pathway pair, we determined the number of direct interactions between them and compared this value to the one obtained from randomized versions of the KEGG signaling network to calculate z-scores and empirical p-values, as described in the previous section. We finally ranked pathway pairs by their empirical p-values and z-scores, prioritizing pairs with the lowest p-values and highest z-scores as the most likely to crosstalk.

### Shortest paths between signaling pathways (Crosstalk Statistic χ)

Following the procedure in (16), we used the shortest paths between the membrane receptor proteins (receptors) and the transcription factors (TFs) in each KEGG signaling pathway to calculate the crosstalk statistic χ. The receptor and TF data were obtained from Almen et al. (81) and Lambert et al. (82), respectively. For the receptor data, we parsed gene symbols from International Protein Index (IPI) Descriptions. We converted both sets of gene symbols to Entrez IDs. We extracted K shortest paths between receptors and TFs using Yen’s algorithm (83) to calculate the crosstalk statistic χ as described in (16) for K values between 1 and 100. We repeated the χ calculation on randomized networks to obtain z-scores and empirical p-values. We then ranked pathway pairs by their empirical p-values and z-scores, prioritizing pairs with the lowest p-values and highest z-scores as the most likely to crosstalk.

To compare the predictive performance of the above methods, we used their respective ranked list of pathway pairs to generate Receiver Operating Characteristic (ROC) and Precision-Recall (PR) curves, and calculated the area under these curves. Since all methods have instances where the magnitude of the respective crosstalk metric for a pair of pathways cannot be determined (e.g. due to the absence of node or edge overlap between pathways, the absence of TFs or receptors in a given pathway or shortest paths connecting them, the lack of statistically over-represented multilink types), we adopted two strategies, one deterministic and one stochastic, when generating the ROC and PR curves. For the deterministic assessment, we only used the pathway pairs that were “detected” (i.e., pathway pairs that were assigned crosstalk statistics and corresponding p-values and z-scores and that could hence be ranked) in the ROC and PR curves. For the stochastic assessment, we used all 600 pathway pairs in the benchmark, including the ones that did not have crosstalk metrics, and shuffled the ranks of such pairs 1,000 times. This procedure resulted in ROC and PR curves that were “fuzzy” after the detection threshold, and the area under the curves (AUCs) (Supplementary Figure 11) for the stochastic approach were represented by the mean and standard deviation of all shuffled AUCs.

### Using PubMed queries to assess the discovery predictions

Our complete set of 60 KEGG signaling pathways encompasses 3,540 ordered pathway pairs whose crosstalk can be explored. We call the pathway pairs outside of the 600 benchmark pairs our “discovery” set. Since there are currently no literature-curated databases on crosstalk among this larger set, we devised a complementary approach that utilizes PubMed queries to look for evidence of crosstalk among the predictions made by MuXTalk on the discovery set. Using the rentrez R package (84), which leverages the Medical Subject Heading (MeSH) term search functionality of PubMed for a controlled vocabulary of medical terms that allows more clinically-informed queries. We manually curated a list of keywords for each signaling pathway (Supplementary Table 6). We then performed two versions of automated PubMed queries, following an approach similar to (37), as follows: Query 1 (more stringent): (Pathway A) AND (Pathway B) AND (signaling) AND (crosstalk) and Query 2 (less stringent): (Pathway A) AND (Pathway B) AND (signaling) AND (pathway). We considered each query that resulted in at least one publication for the given pathway pair as a potential positive and created precision-rank plots for the top-ranked pathway pairs to track MuXTalk’s rate of capturing these potential positives.

## Supplementary Figures and Tables

**Supplementary Figure 1:**
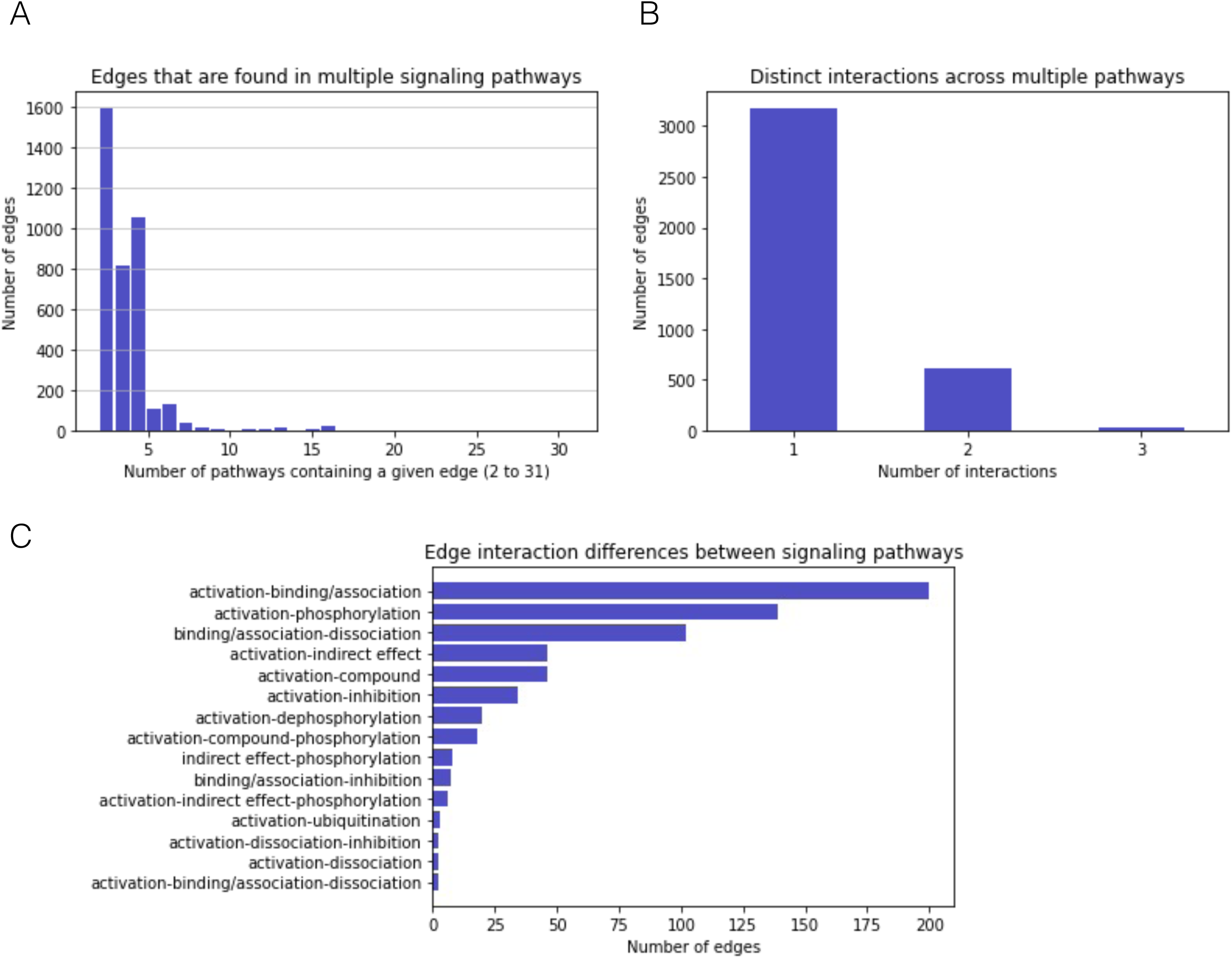
**(A)** Distribution of the number of pathways of which each edge is a member. **(B)** The disctribution of the number of distinct interaction types in cases where the edge belongs to multiple pathways. **(C)** The breakdown of distinct interaction types in cases where the edge represents multiple distinct types of interactions across multiple pathways.

**Supplementary Figure 2:**
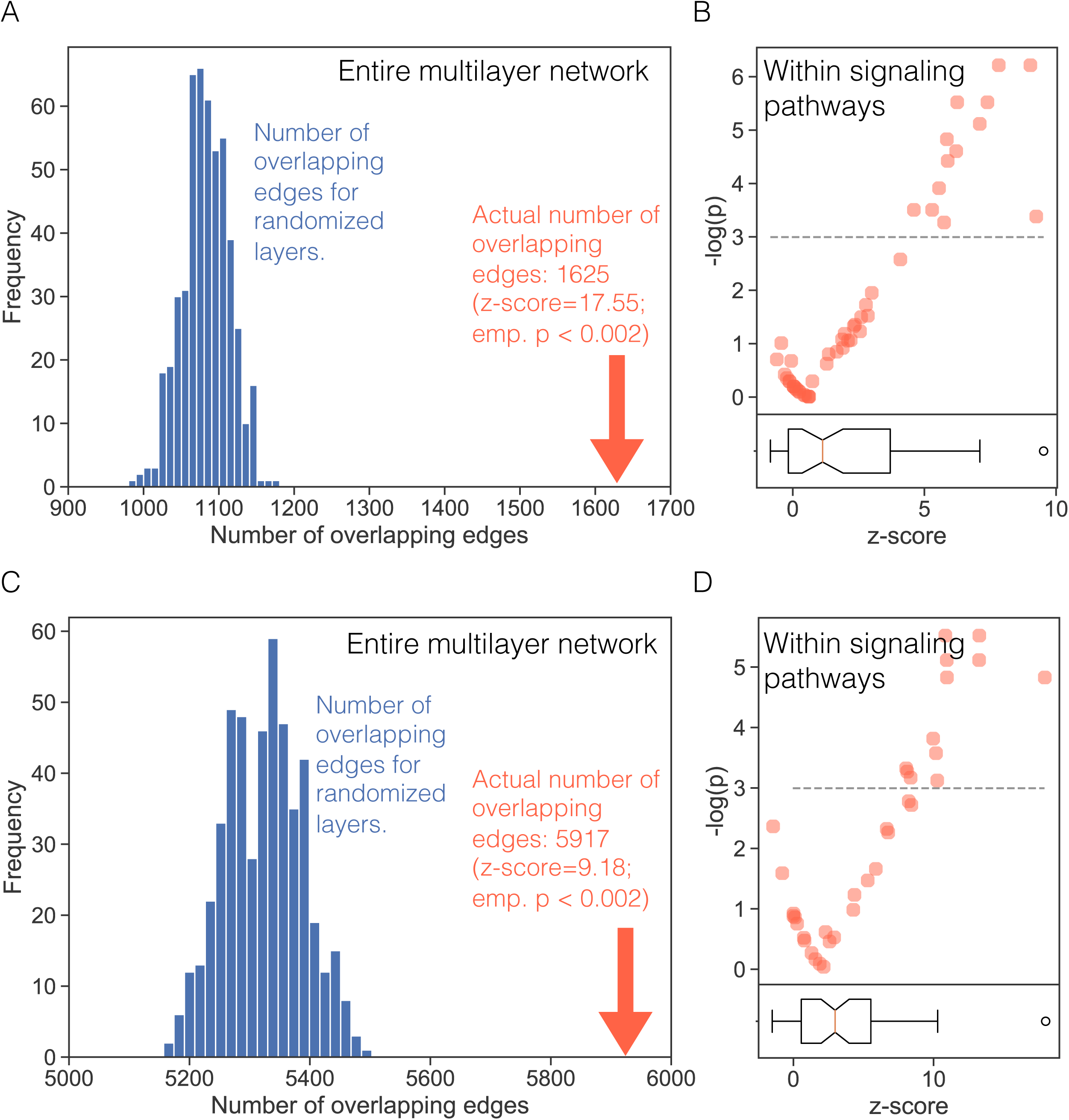
**(A, C)** The number of overlapping signaling and regulatory edges in the multilayer network with a GRN layer of p-value threshold (p<10-5) and (p<10-4), respectively. The blue bars show the distribution of overlapping edges for the randomized networks and the red arrow indicates the overlap for the actual multilayer network. **(B, D)** The -log(empirical p) values for overlap within each signaling pathway in the multilayer network with a GRN layer of p-value threshold (p<10-5) and (p<10-4), respectively. Each dot represents a KEGG pathway. The boxplot indicates the distribution of z-scores for overlap within each pathway.

**Supplementary Figure 3:**
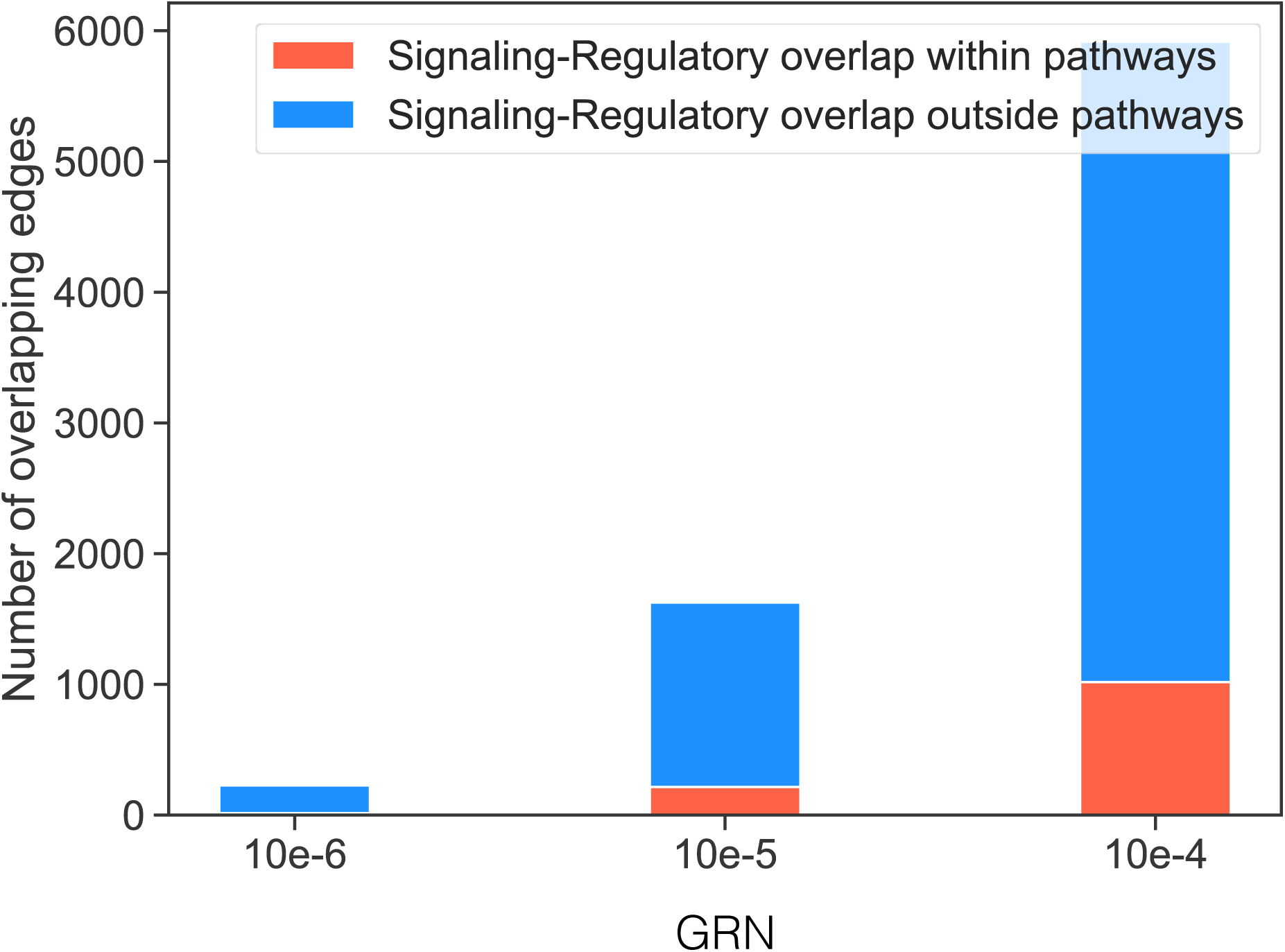
The number of overlapping edges between the signaling and the regulatory layer for GRNs with different p-value thresholds. Blue bars indicate the proportion of overlap outside, or between, KEGG signaling pathways and the red bars indicate the proportion within KEGGG signaling pathways.

**Supplementary Figure 4:**
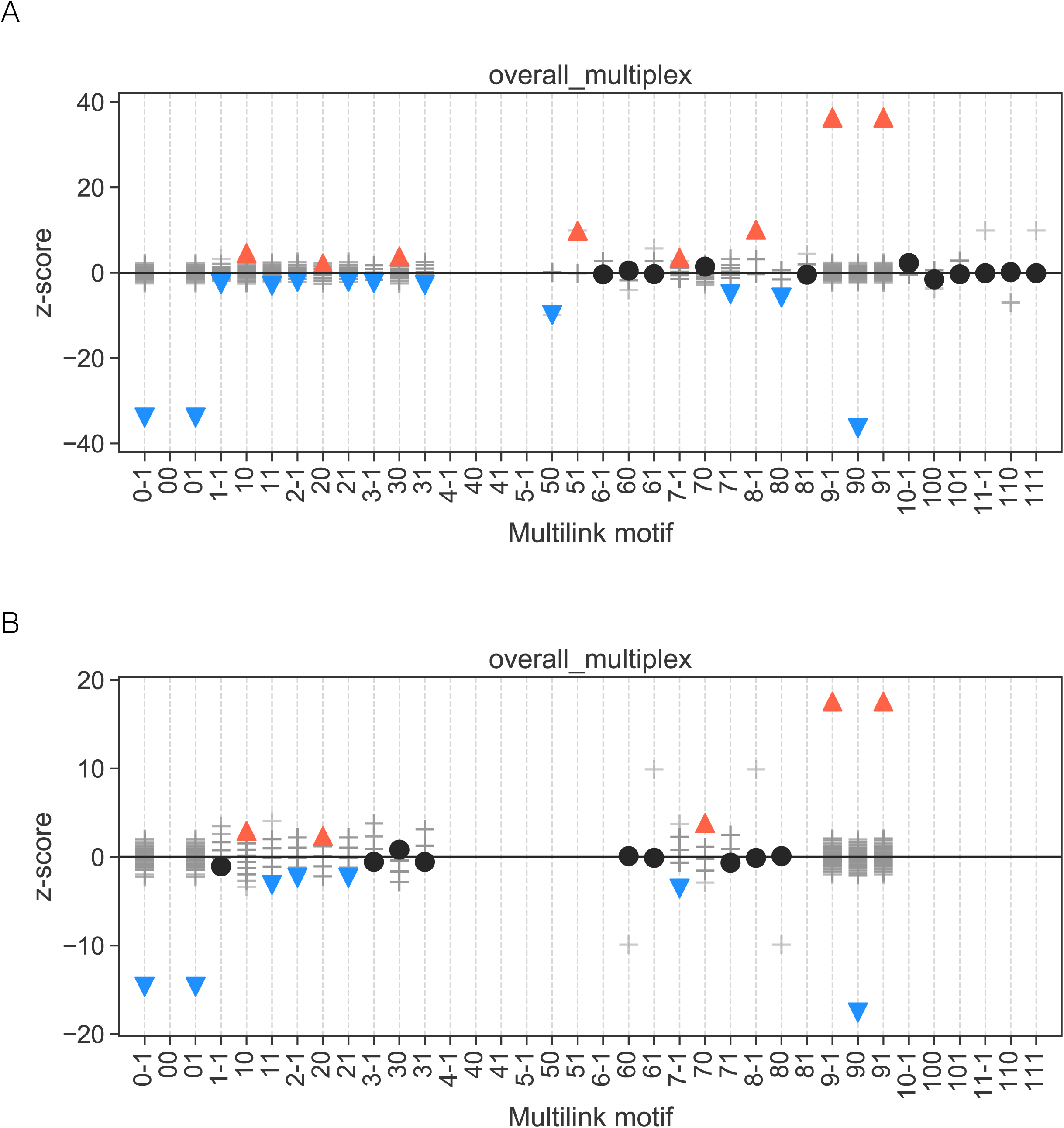
Multilink profiles of all edges in the multilayer network with a GRN layer of p-value threshold (p<10-5) **(A)** and (p<10-6) **(B)**. Multilink types are represented by numerical indicators where the first one or two digits represent the signaling edge type and the last digit represents the regulatory interaction direction. Red triangles pointing up and blue triangles pointing down indicate statistically over-represented (z>0; emp. p≤0.05) and under-represented (z<0; emp. p≤0.05) multilink types, respectively. Black circles denote statistically insignificant multilink types (emp. p>0.05).

**Supplementary Figure 8:**
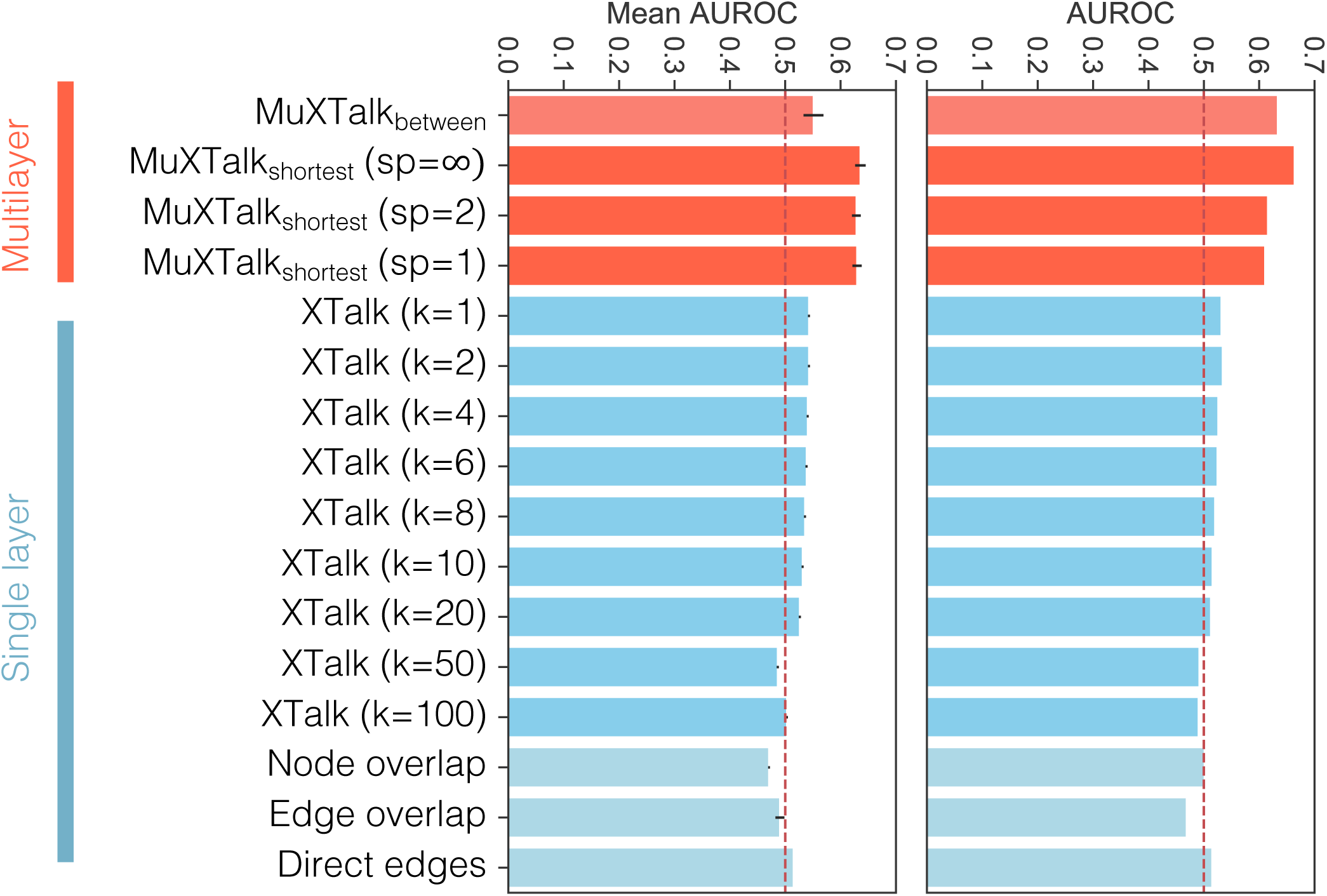
Area under the receiver operating characteristic (AUROC) curves for MuXTalk (red) and four other methods (blue) for the stochastic (left) and deterministic (right) versions of the benchmark. Error bars indicate the standard deviation. MuXTalk was run on the multilayer network with a GRN layer of p-value threshold (p<10-6)

**Supplementary Figure 9:**
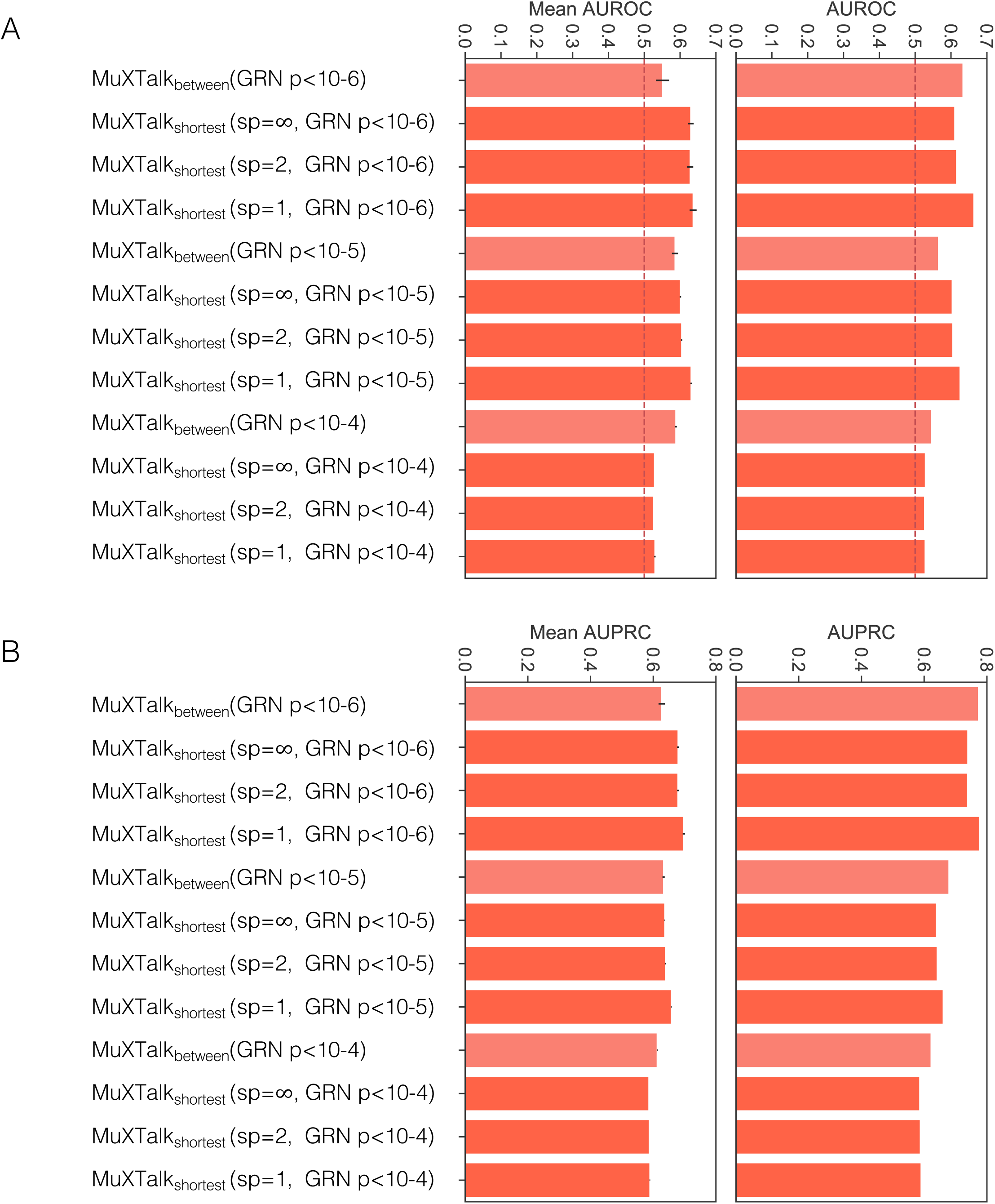
Area under the receiver operating characteristic (AUROC) **(A)** and precision-recall curves (AUPRC) **(B)** for MuXTalk for the stochastic (left) and deterministic (right) versions of the benchmark, for multilayer networks with GRN layers of p-value threshold (p<10-6, p<10-5 and p<10-4). Error bars indicate the standard deviation.

**Supplementary Figure 10:**
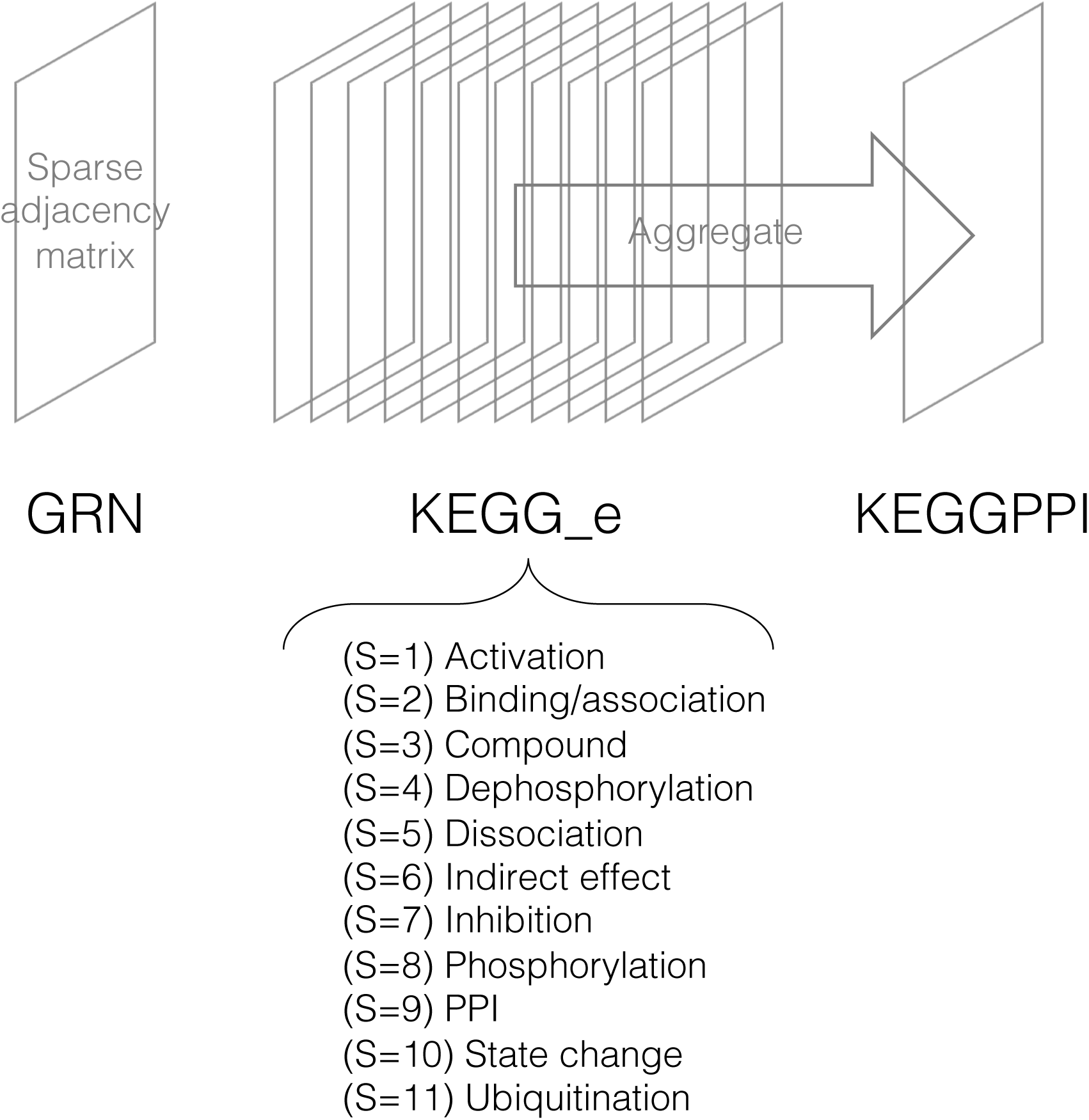
Schematic showing the sparse adjacency matrices used in MuXTalk. For each type of sparse matrices, we generated ensembles (N=500) of randomized versions on which to calculate z-scores and empirical p-values.

**Supplementary Figure 11:**
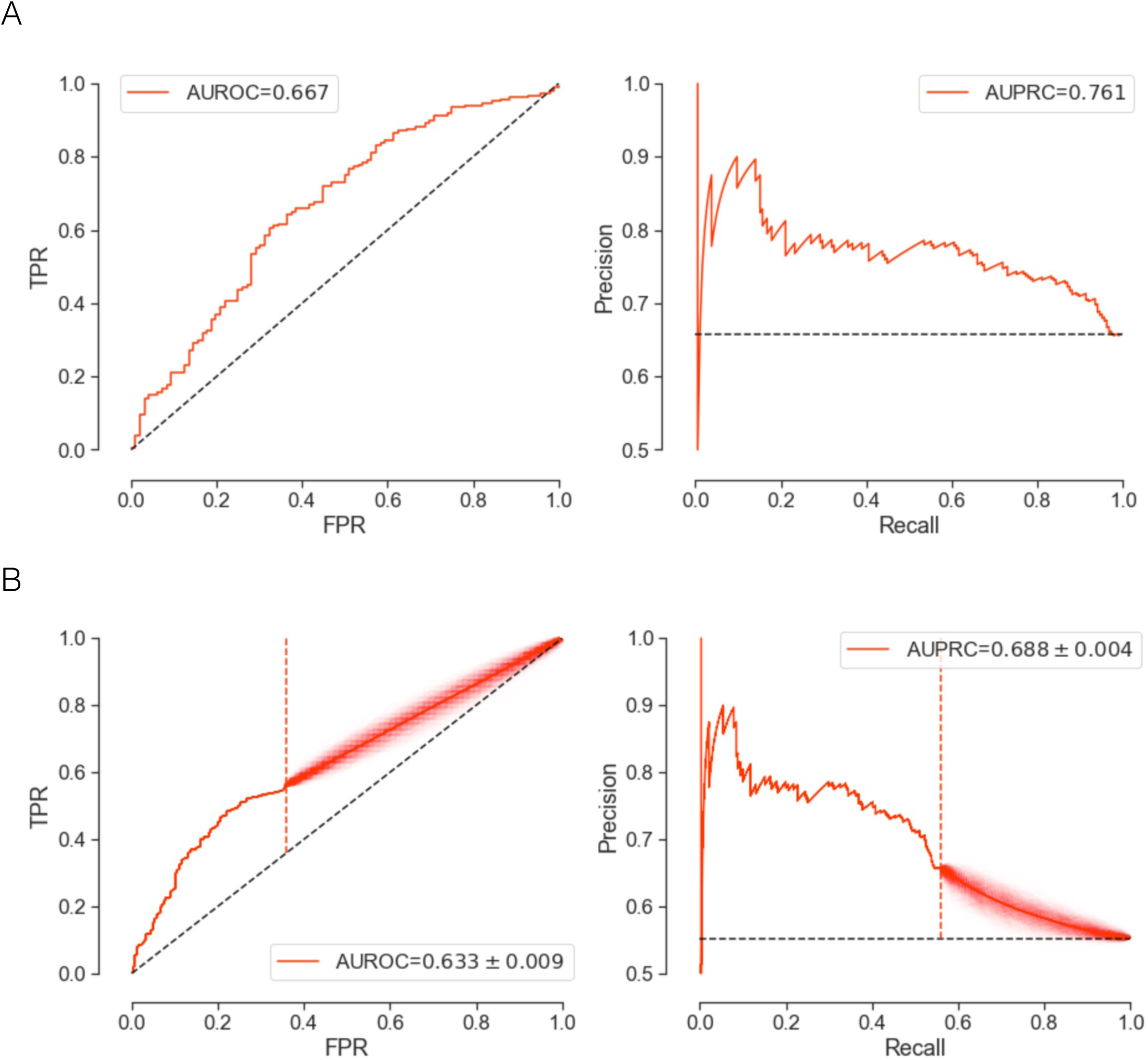
Examples showing the deterministic **(A)** and stochastic **(B)** versions of the benchmark.

**Supplementary Table 1.**

| KEGG Signaling Pathway | n_nodes | n_edges | density | benchmark |
| --- | --- | --- | --- | --- |
| Adherens junction | 68 | 207 | 0.04543459 | Y |
| Adipocytokine signaling pathway | 61 | 220 | 0.06010929 | Y |
| Adrenergic signaling in cardiomyocytes | 150 | 880 | 0.0393736 | N |
| AGE-RAGE signaling pathway in diabetic complications | 69 | 234 | 0.04987212 | N |
| AMPK signaling pathway | 86 | 280 | 0.03830369 | N |
| Apelin signaling pathway | 109 | 804 | 0.06829766 | N |
| Apoptosis | 121 | 301 | 0.02073003 | Y |
| B cell receptor signaling pathway | 77 | 201 | 0.03434723 | N |
| C-type lectin receptor signaling pathway | 85 | 188 | 0.02633053 | N |
| Calcium signaling pathway | 152 | 872 | 0.03799233 | N |
| cAMP signaling pathway | 192 | 956 | 0.02606894 | N |
| Cell adhesion molecules (CAMs) | 130 | 634 | 0.03780561 | N |
| cGMP-PKG signaling pathway | 147 | 383 | 0.01784549 | N |
| Chemokine signaling pathway | 187 | 1907 | 0.05482721 | N |
| Cytokine-cytokine receptor interaction | 196 | 257 | 0.00672423 | N |
| ECM-receptor interaction | 77 | 210 | 0.03588517 | N |
| Epithelial cell signaling in Helicobacter pylori infection | 31 | 40 | 0.04301075 | N |
| ErbB signaling pathway | 83 | 212 | 0.03114899 | Y |
| Estrogen signaling pathway | 88 | 231 | 0.03017241 | Y |
| Fc epsilon RI signaling pathway | 58 | 175 | 0.05293406 | N |
| FoxO signaling pathway | 81 | 262 | 0.0404321 | N |
| Glucagon signaling pathway | 76 | 406 | 0.07122807 | N |
| GnRH signaling pathway | 86 | 310 | 0.04240766 | Y |
| Hedgehog signaling pathway | 47 | 196 | 0.0906568 | Y |
| HIF-1 signaling pathway | 61 | 149 | 0.04071038 | Y |
| Hippo signaling pathway | 135 | 676 | 0.03736871 | Y |
| Insulin secretion | 50 | 87 | 0.0355102 | Y |
| Insulin signaling pathway | 127 | 436 | 0.02724659 | Y |
| JAK-STAT signaling pathway | 140 | 3197 | 0.16428571 | Y |
| MAPK signaling pathway | 294 | 2191 | 0.02543475 | Y |
| Melanogenesis | 99 | 446 | 0.0459699 | Y |
| mTOR signaling pathway | 133 | 398 | 0.02267031 | Y |
| Neuroactive ligand-receptor interaction | 190 | 228 | 0.00634921 | N |
| Neurotrophin signaling pathway | 115 | 477 | 0.03638444 | Y |
| NF-kappa B signaling pathway | 65 | 102 | 0.02451923 | Y |
| NOD-like receptor signaling pathway | 148 | 335 | 0.01539805 | N |
| Notch signaling pathway | 47 | 190 | 0.08788159 | Y |
| Oxytocin signaling pathway | 132 | 396 | 0.02290076 | N |
| p53 signaling pathway | 35 | 56 | 0.04705882 | N |
| Phosphatidylinositol signaling system | 86 | 2747 | 0.37578659 | N |
| Phospholipase D signaling pathway | 112 | 450 | 0.03619691 | N |
| PI3K-Akt signaling pathway | 265 | 2936 | 0.04196684 | N |
| PPAR signaling pathway | 6 | 24 | 0.8 | N |
| Prolactin signaling pathway | 59 | 127 | 0.0371128 | Y |
| Rap1 signaling pathway | 204 | 1227 | 0.02962909 | N |
| Ras signaling pathway | 226 | 1586 | 0.03118977 | N |
| Relaxin signaling pathway | 109 | 573 | 0.04867482 | N |
| Retrograde endocannabinoid signaling | 103 | 672 | 0.06396345 | N |
| RIG-I-like receptor signaling pathway | 47 | 85 | 0.03931545 | N |
| Signaling pathways regulating pluripotency of stem cells | 98 | 429 | 0.04512939 | N |
| Sphingolipid signaling pathway | 94 | 261 | 0.02985587 | N |
| T cell receptor signaling pathway | 94 | 262 | 0.02997026 | N |
| TGF-beta signaling pathway | 86 | 302 | 0.04131327 | Y |
| Thyroid hormone signaling pathway | 82 | 382 | 0.0575128 | N |
| Thyroid hormone synthesis | 44 | 103 | 0.05443975 | Y |
| TNF signaling pathway | 72 | 143 | 0.0279734 | Y |
| Toll-like receptor signaling pathway | 69 | 136 | 0.02898551 | Y |
| VEGF signaling pathway | 58 | 227 | 0.06866304 | Y |
| Viral protein interaction with cytokine and cytokine receptor | 90 | 101 | 0.01260924 | N |
| Wnt signaling pathway | 146 | 1119 | 0.05285782 | Y |

**Supplementary Table 2.**

| GRN | # TFs | # Targets | # Nodes | # Edges | Density |
| --- | --- | --- | --- | --- | --- |
| 10e-6 | 468 | 10,063 | 10,245 | 34,643 | 0.00066 |
| 10e-5 | 605 | 11,513 | 11,708 | 348,664 | 0.00509 |
| 10e-4 | 632 | 11,517 | 11,716 | 1,969,966 | 0.02870 |

**Supplementary Table 3.**

| Analysis | Randomized multilayer ensemble index | Notes |
| --- | --- | --- |
| Overall multilink statistics | 0-99 | 100 randomizations of the entire multilayer network |
| Pathway-specific multilink statistics | 100-6199 | 61x100 randomizations of the entire multilayer network |
| MuXTalk<sub>between</sub> benchmark | 6200-66199 | 600 (# of pathway pairs in the benchmark)x100 randomizations of the entire multilayer network |
| MuXTalk<sub>shortest</sub> benchmark | 66200-126199 | 600 (# of pathway pairs in the benchmark)x100 randomizations of the entire multilayer network |

**Supplementary Table 4.**
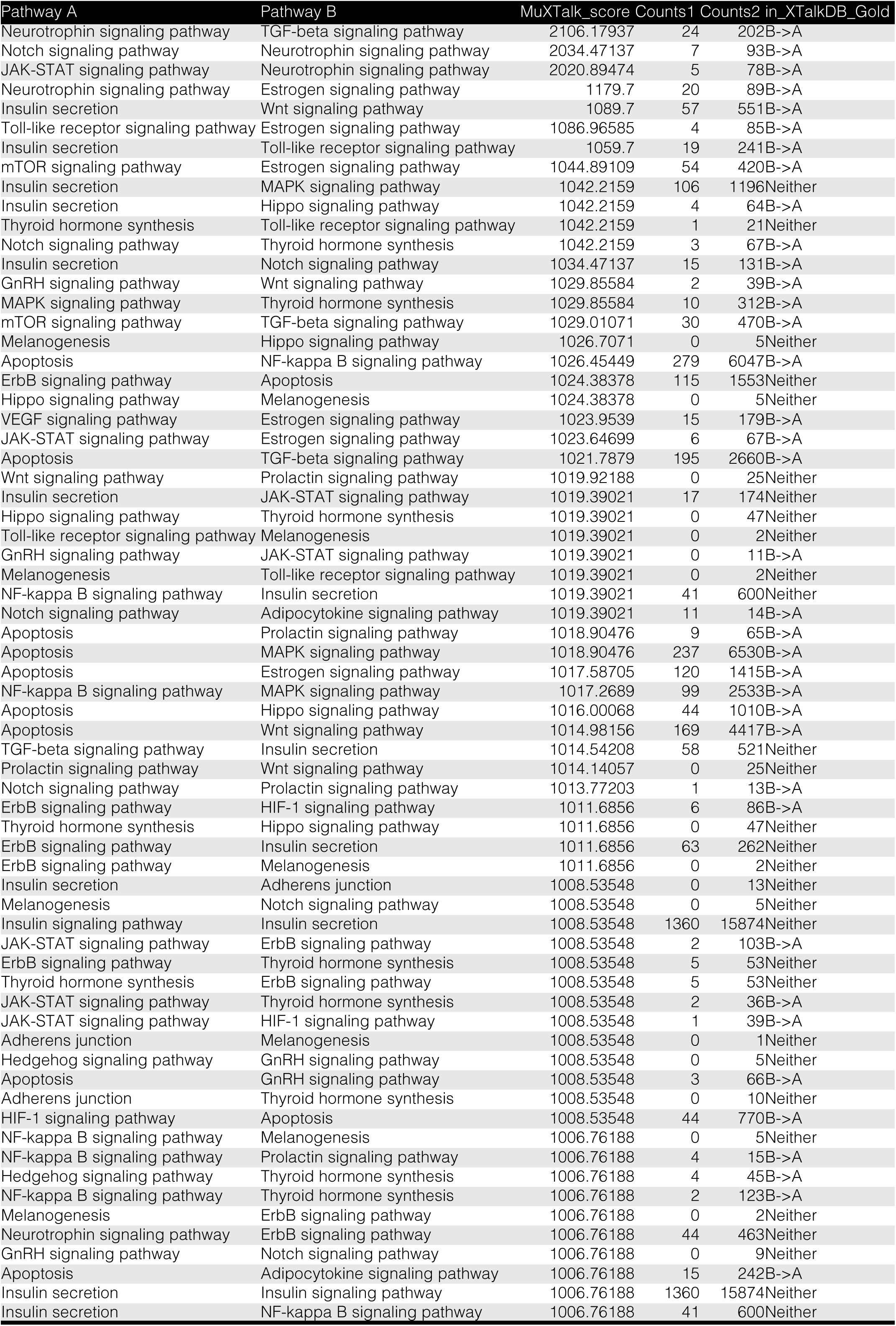
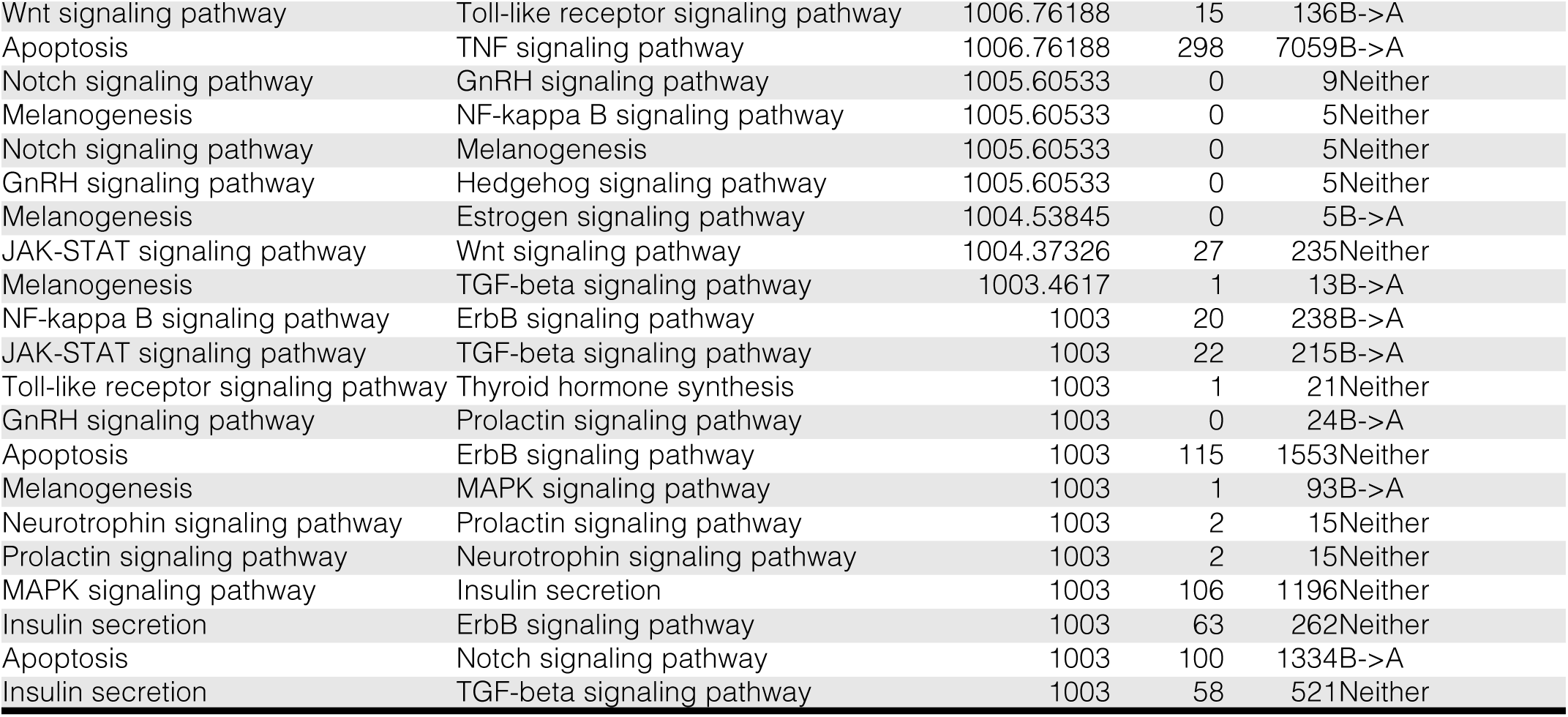

**Supplementary Table 5.**

| Pathway A | Pathway B | MuXTalk_score | Counts1 | Counts2 | in_XTalkDB_Gold |
| --- | --- | --- | --- | --- | --- |
| Apoptosis | Estrogen signaling pathway | 2018.83733 | 120 | 1415 | B->A |
| Neurotrophin signaling pathway | TGF-beta signaling pathway | 2015.85896 | 24 | 202 | B->A |
| Apoptosis | TGF-beta signaling pathway | 2014.77281 | 195 | 2660 | B->A |
| mTOR signaling pathway | TGF-beta signaling pathway | 1022.12982 | 30 | 470 | B->A |
| mTOR signaling pathway | Estrogen signaling pathway | 1016.5687 | 54 | 420 | B->A |
| Apoptosis | Hippo signaling pathway | 1014.45492 | 44 | 1010 | B->A |
| Apoptosis | Wnt signaling pathway | 1014.30758 | 169 | 4417 | B->A |
| Wnt signaling pathway | Prolactin signaling pathway | 1014.07395 | 0 | 25 | Neither |
| Apoptosis | Melanogenesis | 1013.22843 | 2 | 42 | B->A |
| Melanogenesis | Prolactin signaling pathway | 1013.22175 | 0 | 1 | Neither |
| Prolactin signaling pathway | Melanogenesis | 1013.10225 | 0 | 1 | Neither |
| TNF signaling pathway | Melanogenesis | 1013.02262 | 0 | 3 | B->A |
| VEGF signaling pathway | Estrogen signaling pathway | 1012.83946 | 15 | 179 | B->A |
| Toll-like receptor signaling pathway | ErbB signaling pathway | 1012.82627 | 3 | 62 | Neither |
| Neurotrophin signaling pathway | Estrogen signaling pathway | 1012.48041 | 20 | 89 | B->A |
| Prolactin signaling pathway | Wnt signaling pathway | 1012.11132 | 0 | 25 | Neither |
| Adherens junction | TNF signaling pathway | 1011.56673 | 2 | 28 | B->A |
| ErbB signaling pathway | Toll-like receptor signaling pathway | 1009.9889 | 3 | 62 | Neither |
| Insulin secretion | TGF-beta signaling pathway | 1009.64181 | 58 | 521 | Neither |
| JAK-STAT signaling pathway | TGF-beta signaling pathway | 1009.15545 | 22 | 215 | B->A |
| Adherens junction | Prolactin signaling pathway | 1008.45344 | 0 | 1 | Neither |
| Prolactin signaling pathway | Adherens junction | 1007.98306 | 0 | 1 | Neither |
| Melanogenesis | TGF-beta signaling pathway | 1006.98922 | 1 | 13 | B->A |
| Apoptosis | Insulin secretion | 1006.76188 | 124 | 2252 | B->A |
| TGF-beta signaling pathway | Insulin secretion | 1006.34269 | 58 | 521 | Neither |
| Adherens junction | VEGF signaling pathway | 1005.26494 | 0 | 28 | B->A |
| Prolactin signaling pathway | Hippo signaling pathway | 1004.68472 | 0 | 3 | Neither |
| Melanogenesis | GnRH signaling pathway | 1004.61921 | 0 | 0 | Neither |
| Adherens junction | HIF-1 signaling pathway | 1004.28279 | 0 | 3 | B->A |
| Hippo signaling pathway | Prolactin signaling pathway | 1004.26348 | 0 | 3 | Neither |

**Supplementary Table 6.**
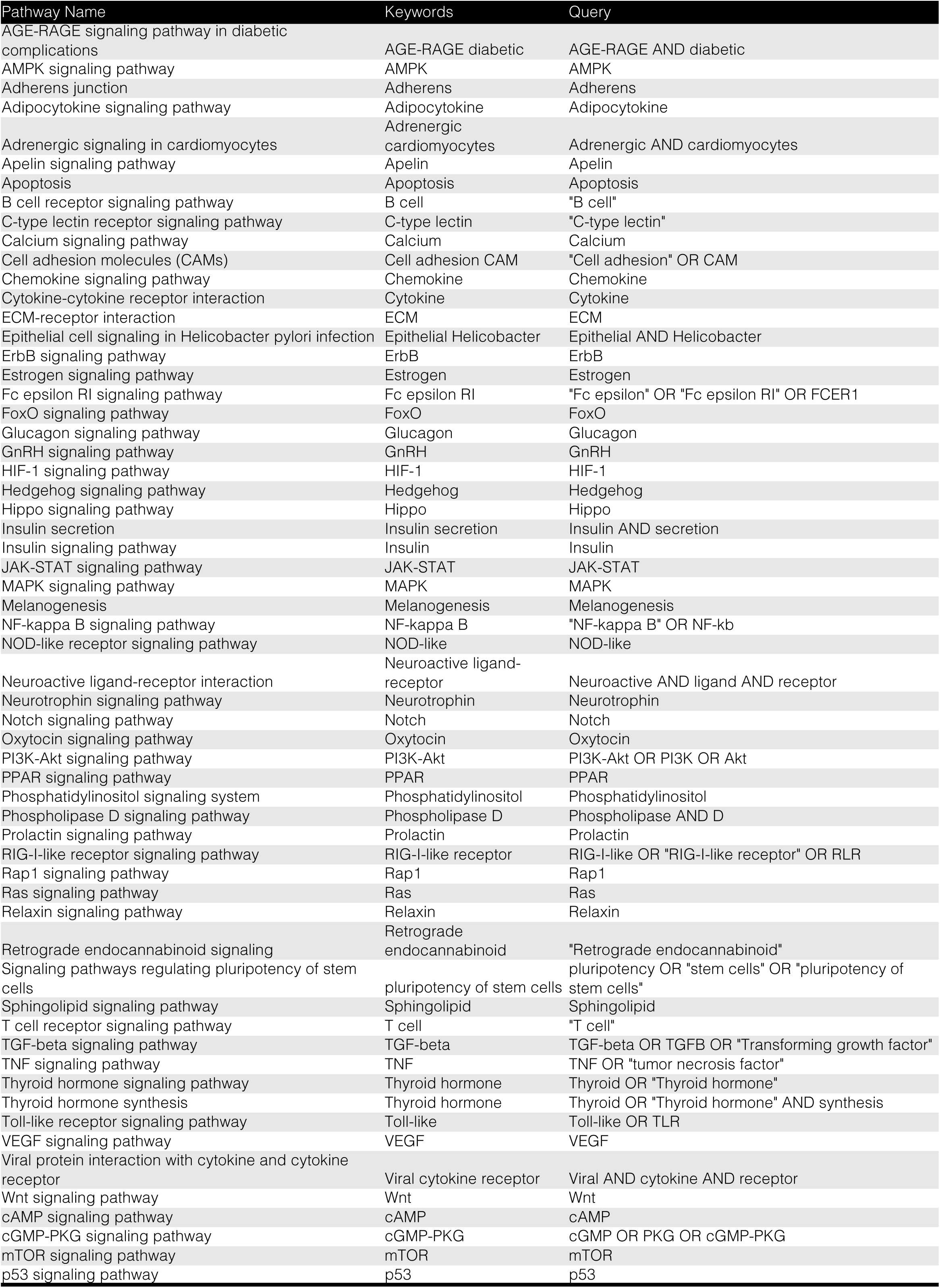

