## Supplementary figures and images for "Detecting and dissecting signaling crosstalk via the multilayer network integration of signaling and regulatory interactions"

### Supplementary Figure 4

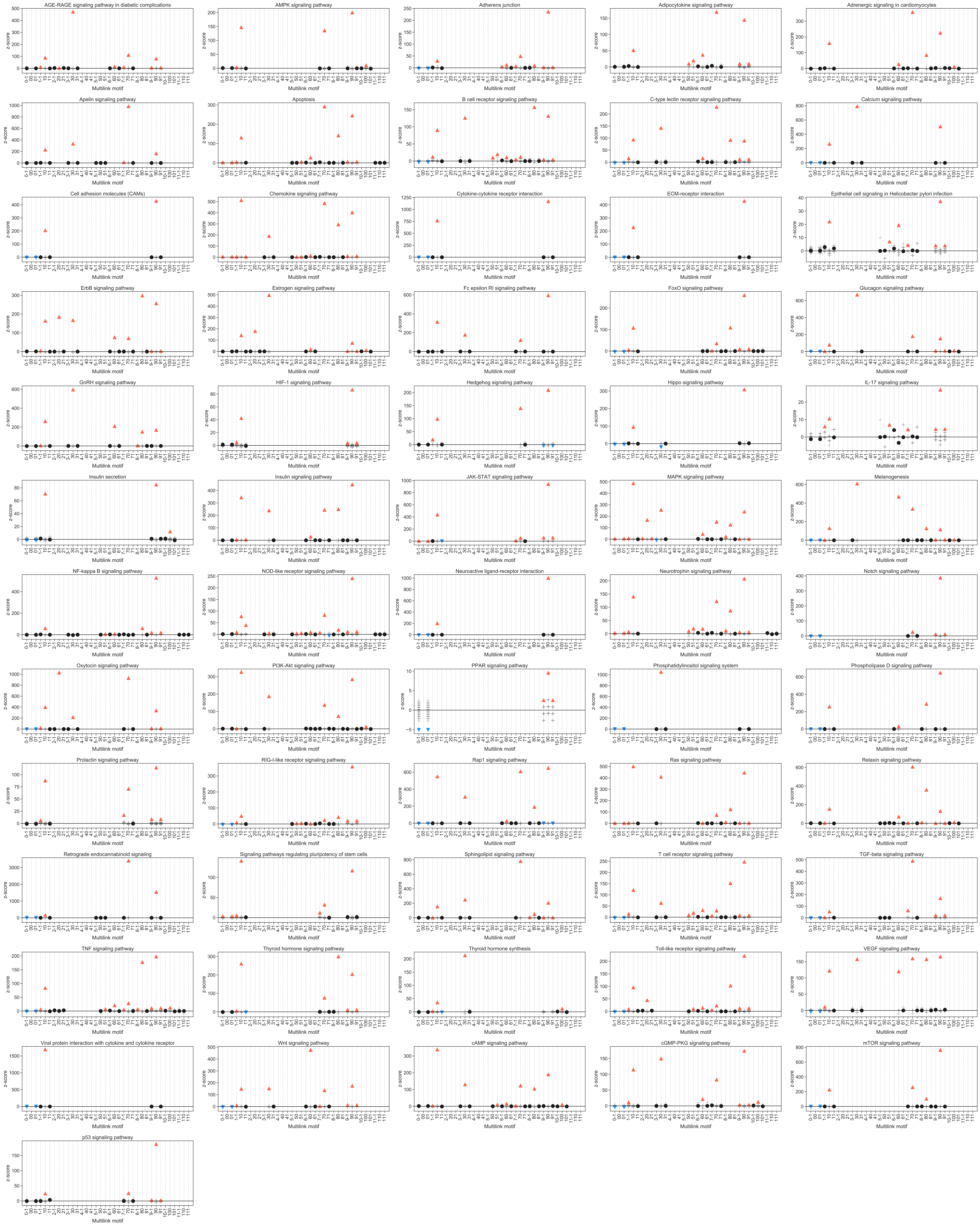

### Supplementary Figure 5

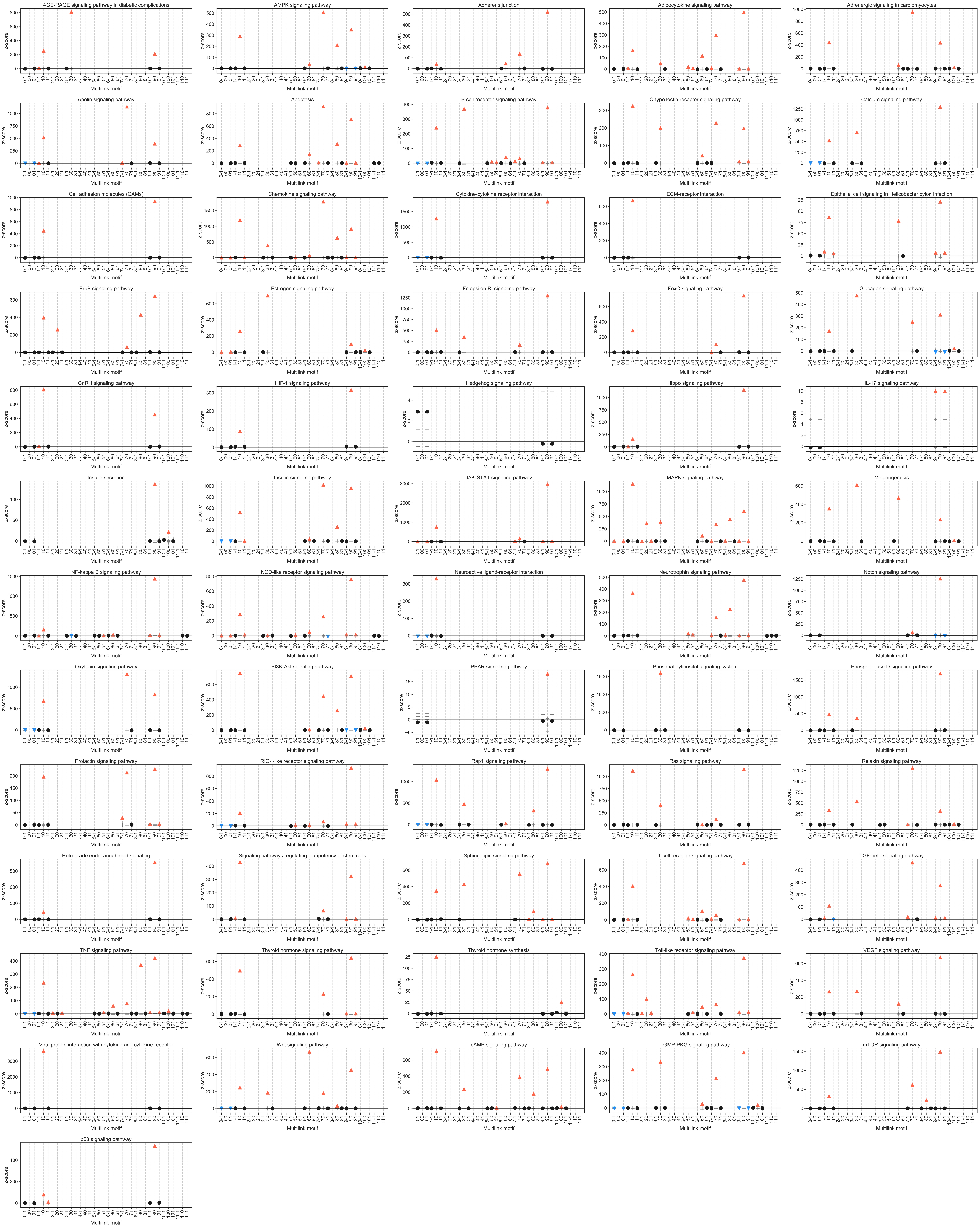

### Supplementary Figure 6

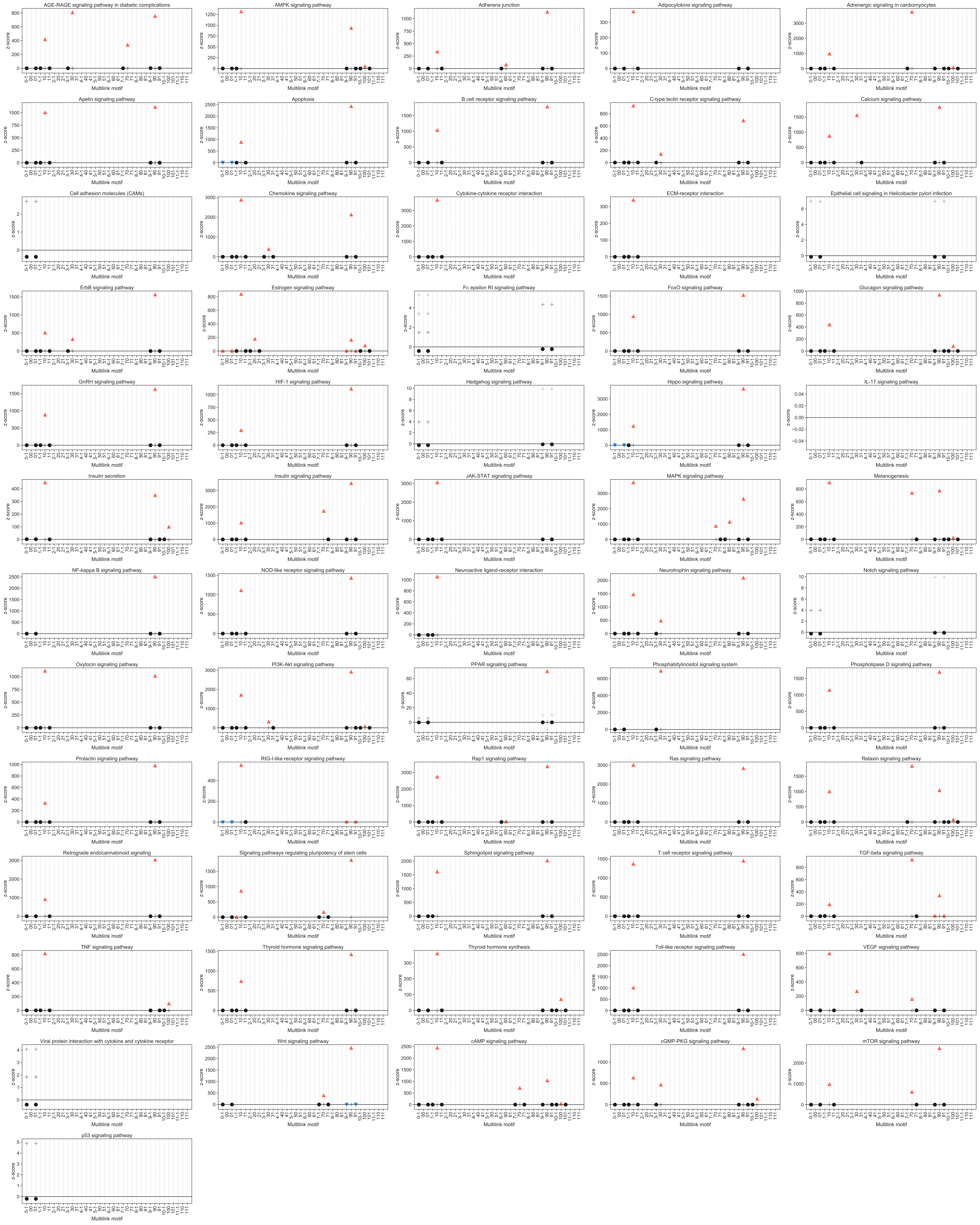
